# Local Field Potentials in a Pre-motor Region Predict Learned Vocal Sequences

**DOI:** 10.1101/2020.06.30.179861

**Authors:** Daril E. Brown, Jairo I. Chavez, Derek H. Nguyen, Adam Kadwory, Bradley Voytek, Ezequiel Arneodo, Timothy Q. Gentner, Vikash Gilja

## Abstract

Neuronal activity within the premotor region HVC is tightly synchronized to, and crucial for, the articulate production of learned song in birds. Characterizations of this neural activity typically focuses on patterns of sequential bursting in small carefully identified subsets of single neurons in the HVC population. Much less is known about population dynamics beyond the scale of individual neurons. There is a rich history of using local field potentials (LFP), to extract information about behavior that extends beyond the contribution of individual cells. These signals have the advantage of being stable over longer periods of time and have been used to study and decode complex motor behaviors, such as human speech. Here we characterize LFP signals in the putative HVC of freely behaving male zebra finches during song production, to determine if population activity may yield similar insights into the mechanisms underlying complex motor-vocal behavior. Following an initial observation that structured changes in the LFP were distinct to all vocalizations during song, we show that it is possible to extract time varying features from multiple frequency bands to decode the identity of specific vocalization elements (syllables) and to predict their temporal onsets within the motif. This demonstrates that LFP is a useful signal for studying motor control in songbirds. Surprisingly, the time frequency structure of putative HVC LFP is qualitatively similar to well established oscillations found in both human and non-human mammalian motor areas. This physiological similarity, despite distinct anatomical structures, may give insight to common computational principles for learning and/or generating complex motor-vocal behaviors.

**Author Summary:** Vocalizations, such as speech and song, are a motor process that requires the coordination of several muscle groups receiving instructions from specific brain regions. In songbirds, HVC is a premotor brain region required for singing and it is populated by a set of neurons that fire sparsely during song. How HVC enables song generation is not well understood. Here we describe network activity in putative HVC that precedes the initiation of each vocal element during singing. This network activity can be used to predict both the identity of each vocal element (syllable) and when it will occur during song. In addition, this network activity is similar to activity that has been documented in human, non-human primate, and mammalian premotor regions tied to muscle movements. These similarities add to a growing body of literature that finds parallels between songbirds and humans in respect to the motor control of vocal organs. Given the similarities of the songbird and human motor-vocal systems these results suggest that the songbird model could be leveraged to accelerate the development of clinically translatable speech prosthesis.

## Introduction

Learned vocalizations, such as speech and song, are generated by the complex coordination of multiple muscle groups that control the vocal organs[1–3]. Like other voluntary movements this coordinated action arises from premotor brain regions [4–8] and is prepared prior to the physical initiation of movement [4,9,10]. How these behaviors are encoded during both preparation and generation remains an important topic of ongoing research. Developing an understanding for how the brain encodes complex sequential motor movements carries implications for the development of neuroprosthetics that aim to return or supplement lost motor function. In addition to their therapeutic benefits, such systems will help create new tools for examining the brain’s mechanisms for learning and executing motor sequences.

At present, studying the motor encoding of speech and other complex motor movements in humans is difficult due to the intrinsic hurdles of doing invasive human clinical studies [11–15] and the complexity of human language [2,14]. Clinical studies in humans are difficult. The clinical setting constrains experimental studies with limits in how long and from where researchers can record data. The study of other motor movements is often supplemented by first studying simpler animal models such as non-human primates [16,17] and rodents [18–21]. These animal models fall short when it comes to more complex freely generated motor sequences such as speech. This is due to none of these models having the ability to learn vocal behavior resembling the complexity of human speech [2,22]. For this reason speech production studies, unlike other motor behavioral fields, have been limited exclusively to invasive [11,23,24] and non-invasive [25] clinical studies in humans. Previous work has found that the same cortical motor neurons that control hand and arm movements are also involved in the production of speech in patients with tetraplegia [26], providing preliminary support that human speech research could also be supplemented by a viable translational animal model. A natural candidate model is the songbird, which is widely used to study complex learned vocal behavior [27,28]. The zebra finch (Taeniopygia guttata), in particular, is known for its precisely timed sequentially structured song which is stable over the course of its adult lifetime. Experiments with this system have yielded insights into the acquisition [29,30], maintenance [31,32] and generation [8,33,34] of complex motor-vocal sequences.

The premotor nucleus HVC, used as a proper name, contains neurons with sparse spiking patterns precisely aligned to vocal production [8,35]. It is considered analogous to premotor cortex in human and non-human primates both functionally [36–39] and genetically [40,41]. HVC provides input to two pathways that lead to neurons within nuclei that control the vocal muscles of the syrinx (Fig 1A). The first is directly through RA in the posterior descending pathway (PDP), which is necessary for both the acquisition and production of learned motor-vocal behavior. For reference to mammalian anatomy, the PDP is analogous to a motor pathway that starts in the cerebral cortex and descends through the brain stem. The second, named the Anterior Forebrain Pathway (AFP), plays a strong role in acquisition only and projects indirectly through several nuclei analogous to a cortical pathway through the basal ganglia and thalamus in mammals [28]. These analogous regions share thousands of convergent genes despite their last shared ancestor being millions of years ago [41]. At a circuit level the similarities between the neural activity in humans and songbirds is much harder to compare. Single unit work and techniques, like those done in songbirds, are difficult to do within a clinical setting. In contrast given the accessibility of the system and a focus on fundamental physiology, songbird studies have largely focused on carefully identified neural units and their spiking activity, with limited examination of other neural features that may correlate with vocalization behavior. Examining neural features that can be recorded from both species and their relationship to motor-vocal behavior will enable direct comparison between the neural activity in birdsong and human speech production. Clarifying whatever similarities (and differences) that may exist and potentially bridging the gap between the two.

**Fig 1.**
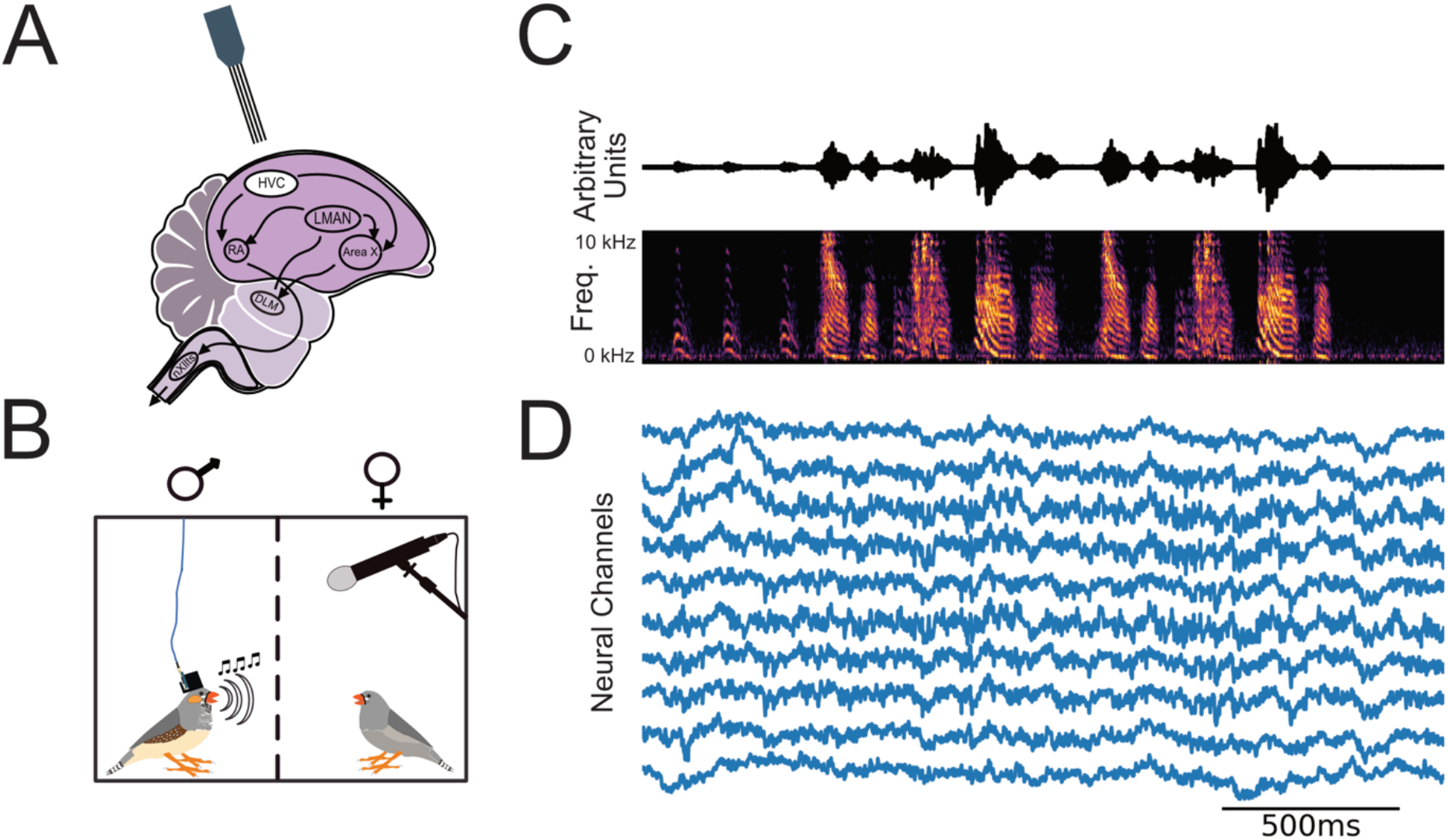
Continuous electrical and audio recording of chronically implanted free behaving male zebra finch (Methods). (A) Four-Shank multi-site Si-Probes were chronically implanted targeting nucleus HVC. HVC provides input to two motor pathways that innervate neurons in the tracheosyringeal half of the hypoglosssal nucleus (nXIIts) that projects to vocal muscles. The posterior descending pathway (PDP) comprises a direct projection to the robust nucleus of the arcopallium (RA) and is necessary for both the acquisition and production of learned vocal behavior (song). The Anterior Forebrain Pathway (AFP), provides an indirect pathway to RA, through Area X, the dorsolateral anterior thalamic nucleus (DLM), and LMAN, and is necessary for song acquisition. The PDP is homologous to a motor pathway in mammals that starts in the cerebral cortex and descends through the brain stem, while the AFP is homologous to a cortical pathway through the basal ganglia and thalamus. (B) Neural and audio recording apparatus. We recorded LFP and vocal activity from male zebra finches stimulated to sing by the presence of a female conspecific. (C) Exemplar sound pressure waveform (1.3 seconds in duration, top) from bird z007 above the corresponding spectrogram. (D) Voltage traces (μV) of ten randomly selected channels of simultaneously recorded neural activity aligned to the song during audio recording, after a 400 Hz low pass filter.

Local Field Potentials (LFP) are thought to reflect both aggregate local postsynaptic activity and presynaptic inputs to the recording site [42]. Because stable single units can be difficult to acquire and maintain in humans and non-human primates there is a rich history of literature looking at both spiking and LFP for understanding, decoding and predicting motor production [43–47]. At present most work with HVC in songbirds has focused almost exclusively on single unit spiking activity [6,28,34,48,49], and limited work has focused on the structure of LFP and how they relate to song production. The most detailed characterization of this signal in songbirds being LFP’s relation to interneuron synchrony [33]. This leaves a gap in the literature regarding the structure of LFP activity in HVC and whether its characteristics have any similarities to LFP in human, non-human primates, or mammalian premotor and motor regions.

We addressed this gap by chronically implanting and recording from freely behaving male zebra finch (Fig 1B) and analyzing the LFP in relation to each bird’s performed song (Fig 1C and D). Our results show narrow-band oscillations similar to those reported in human, non-human primate, and rodent motor electrophysiology literature. Further we provide evidence that phase and amplitude modulation within these frequency bands is predictive of vocalization behavior.

## Results

Adult male zebra finches (n=3) were chronically implanted with a lightweight custom 3D printed microdrive (Fig 1B). Local field potentials and vocal behavior were simultaneously recorded using a laminar silicone probe and an omnidirectional microphone, respectively (Fig 1C and 1D) (see Methods). A male zebra finch’s learned song consists of 1-7 repetitions of a motif, each of which is composed of a precisely ordered sequence of 3-10 discrete units called syllables. Song motifs are also grouped into larger structures called bouts, which consist of multiple repetitions of motifs. The bout is typically preceded by a variable number of repetitions of the same note which are called Introductory Notes. Syllables of each bird’s learned song, other non-learned vocalizations, and tokens of non-vocal intervals were segmented and annotated within periods of vocal behavior, referred to as Vocally Active Periods (VAP), using acoustic landmarks (Fig 2). Temporal boundaries of VAPs were used to define behaviorally relevant epochs in the simultaneously recorded neural signal. Subjects were recorded for 1-10 hrs per day, and statistics of recorded behavior are documented in S1 and S2 Table. To maintain statistical power, we selected the two days with the most vocal activity from each bird for the analyses reported here (S3-S5 Table). Results from days with fewer recorded motifs were qualitatively similar.

**Fig 2:**
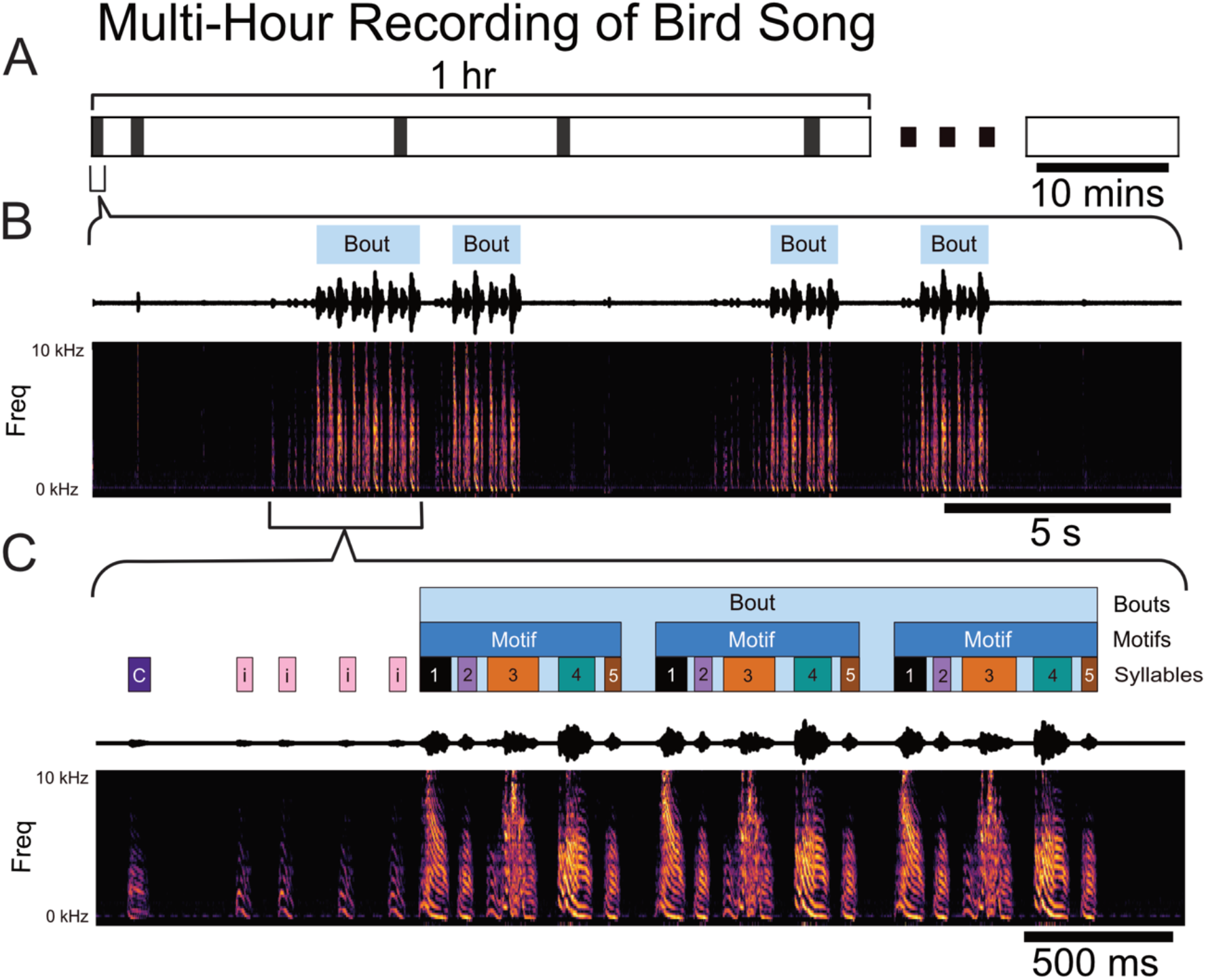
Temporal structure of vocal behavior. (A) Schematic showing the intermittent occurrence of vocally active periods (VAPs, black bar) at sporadic intervals throughout the first hour of a multiple-hours-long continuous recording session of one male zebra finch (z007). (B) Zoomed in view of one 25 second long VAP comprising several bouts, denoted by light blue rectangles above the sound pressure waveform and corresponding spectrogram. (C) Zoomed in view of the first bout in segment (B) showing introductory notes, typically repeated a variable number of times prior to the start of the bout and labeled as ‘i’, other vocalizations not part of the courtship song, labeled ‘C’, and syllables comprising the courtship song, labeled ‘1’, ‘2’, ‘3’, ‘4’, ‘5’, based on their sequential order in the motif.

### Song-related LFP spectral changes in putative HVC

Fig 3A illustrates the spectrotemporal characteristics of the LFP amplitude during the preparation and production of bird song, with an exemplar single trial time-varying power spectral density (PSD) during the course of 3 separate bouts. Each frequency is scaled to its percentage of the mean amplitude during the respective VAP (see Methods). By visual inspection of individual trials we noted a broadband increase in power in all frequencies above roughly 50 Hz prior-to and during vocal production (S2 Fig). This occurs simultaneously with rhythmic changes in frequencies below roughly 50Hz. These changes, which have contributions from both phase and amplitude, were observed to start just prior to song onset and end immediately after vocal production has stopped. Similar patterns emerged in each of the birds and within birds across days (S3 and S4 Fig.). When inspecting single trials changes in LFP amplitude appeared to correspond with each instance of vocal behavior, including non-learned portions such as introductory notes.

**Fig 3:**
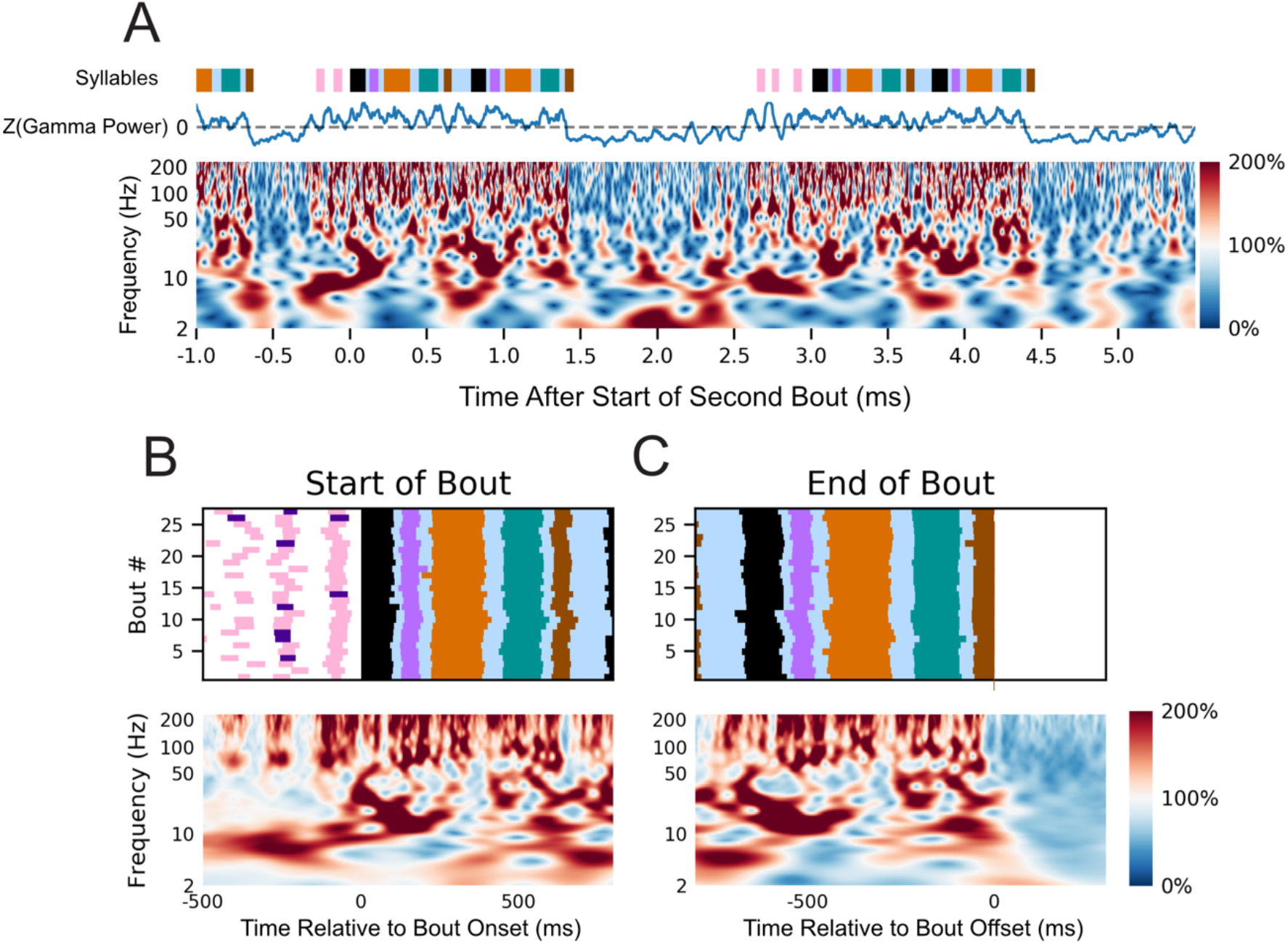
Variation in putative HVC LFP power correlate with vocal behavior. (A) Normalized spectrogram (bottom) of LFP activity from one representative channel (see **Methods**) aligned to the free vocal performance of a male zebra finch (z007) during the production of three separate bouts. Each frequency is scaled to its percentage of the mean amplitude during the respective VAP. Above the spectrogram is the z-scored power of the 50-200 Hz band smoothed with a 50ms rolling mean window aligned to the same behaviors. Colored bars above the spectrum annotate the vocal behavior, with color representing a distinct syllable with the same color coding as Fig 2. (B) Normalized spectrogram (bottom) from the same representative channel as (A) averaged across multiple renditions of similar bouts (n=27, top) aligned to the start of the first motif in the bout. (C) As in (B) but aligned to the end of the last motif in the bout. Above both is a behavioral raster showing the time course of the behavior being averaged (n=27 bouts). No dynamic-time warping was used and all motifs were from the highest yield day for the subject. To ensure the start and end of the bout are unique time periods, only bouts with more than one motif in duration were used. Behaviorally inconsistent bouts were excluded for clarity of visualization, including them does not qualitatively alter the result.

The spectral structure of the LFP activity immediately prior to and during the production of learned song is highly consistent across bouts as shown when they are aligned to their initiation (Fig 3B) and termination (Fig 3C). These results were calculated by aligning to the start of the first syllable of the first motif in the bout (initiation) and the start of the last syllable of the last motif of the bout (termination) for all bouts recorded over the course of a single day. The aligned neural activity was then averaged for each frequency. As with the single trial time-varying PSD, there is an increase in amplitude for all frequencies above roughly 50 Hz (S1 and S2 Fig) and what appeared to be a drop in amplitude for frequencies below roughly 50 Hz. To evaluate the consistency of these changes in amplitude we compared the distributions of power values during song to the distribution of power values during silence using the one-sided z-test. A statistically significant increase in power for frequency bands above 50 Hz was seen across most, if not all, channels for all but one high yield day for z020 (S6 and S7 Table). Changes in amplitude for frequency bands below 50 Hz were more nuanced with no consistent trend across subjects, however frequency changes tended to be consistent across days for two of the birds, particularly z017 and z007 (S6 and S7 Table). Within subject, there was strong stereotyped structure for frequency bands below 50 Hz arising from co-occurrences in changes from both phase and amplitude. S3 and S4 Fig illustrate this stereotypy of time correlated spectral structure in HVC as it relates to the preparation and production of bird song. No dynamic time warping was used to align the behavior.

As in previous work [31,33], we found a strong stereotyped 25-35 Hz oscillation that was consistent across renditions of motifs (Fig 3B and 3C). As shown in Fig 1A, patterned changes in spectrum occur over times scales longer than a single cycle for a given frequency below approximately 50 Hz (S3 Fig and S4 Fig). These oscillations were found to have contributions from both phase and amplitude [33], and occur over several frequencies besides 25-35 Hz. Which led us to ask: what other potential frequencies in the LFP might carry information about the preparation and production of song through either phase or amplitude changes.

### Decoupling LFP power-spectrum reveals song related spectral features

A spectral decomposition technique previously used in human motor cortex was used [50] to determine which frequencies’ amplitude changes were correlated with the production of birdsong. As shown in Fig 4A, on average there was a consistent increase in higher frequencies that are in the range often referred to as “high gamma” [51,52]. The frequency ranges for “High Gamma” vary in the literature but for Miller et. al. it is described as 80-200Hz. Following the steps of the spectral decomposition approach described in the methods, the principal component decomposition of these PSDs found Principal Spectral Components (PSC) similar to those previously reported in [50] (Fig 4 B & C). The PSC’s shown in Fig 4 are results from one channel which was representative of all channels for each subject. The first PSC, or the most significant principal component, in Miller et al is characterized for having its element magnitudes with the same sign being consistently non-zero, and their values being closer to the average across all elements than to zero. This was subsequently shown to reflect a broad spectrum increase in high gamma. In our analysis the most consistent PSC, which is the first principal component, was a spectrum wide-amplitude change that was consistent across channels and subjects (S5 Fig.). The second PSC in Miller et al peaked in the “alpha/low beta range” – alpha is described as 8-12 Hz and beta is described as 12-30 Hz – which mirrors PSC 3 found across subjects and channels in zebra finch (S5 Fig) [50]. These frequency bands in humans have been proposed to reflect the resting rhythms that decrease when cortical areas are activated [53]. The characteristic structure of each PSC were found to be consistent across subjects, channels, and days using the cosine similarity metric (see Methods) (S5 Fig). In addition, this metric also found that each PSC was found to be more similar to one another across subjects than they were with a different PSC (S5 E-G Fig). Although the PSCs were calculated without explicit knowledge of distinct vocally active and inactive time periods, the first three PSCs show that the power spectrum during vocal production is separable from vocally inactive periods (Fig 4C). This suggests that these frequencies could be used to detect the onset of vocal production.

**Fig 4:**
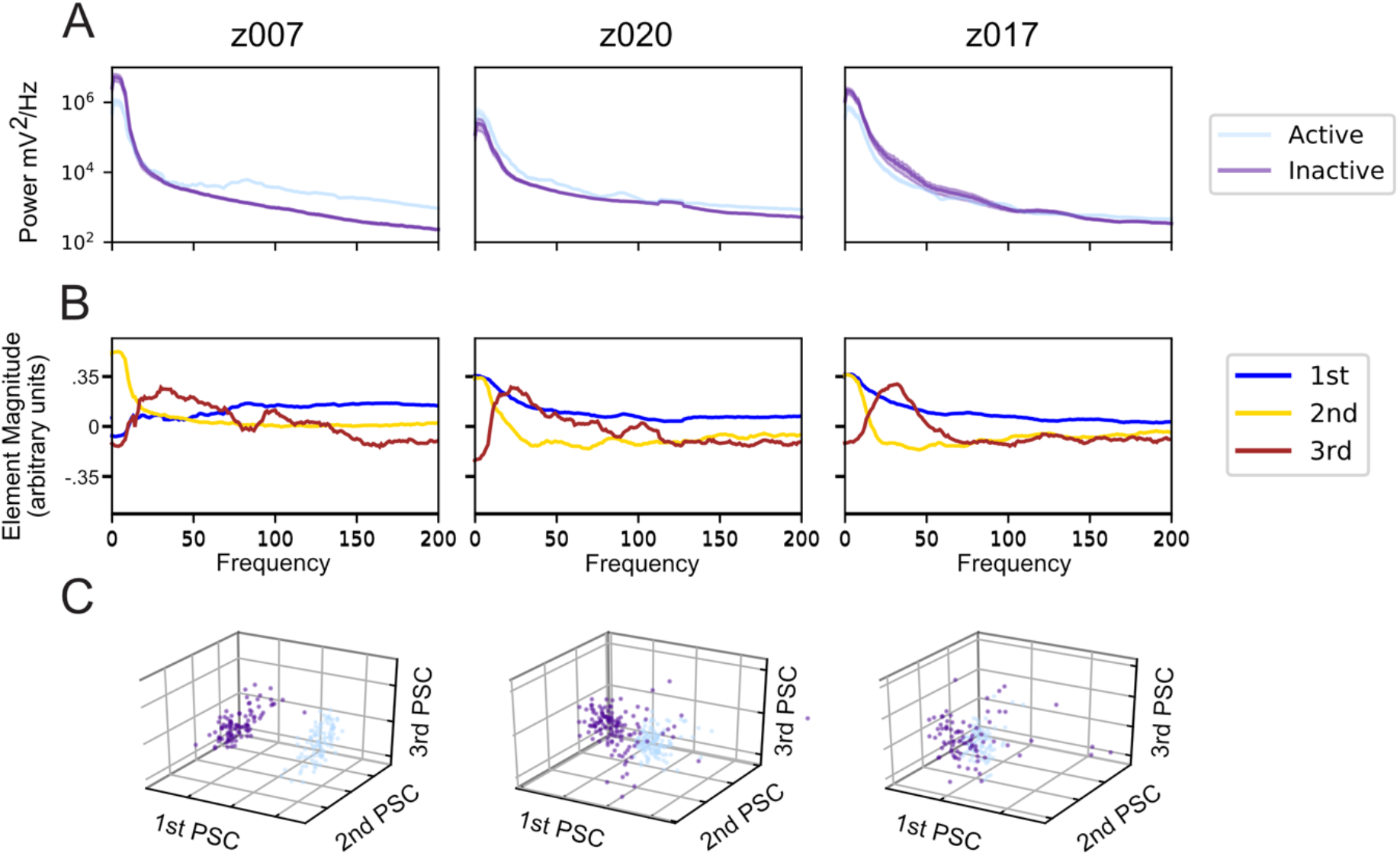
Naïve decomposition of LFP power spectra reveals song correlated features. Representative results from a single electrode channel for each of the three subjects during its Highest Yield Day. (A) Averaged Power spectra during trials centered in 1 second intervals where the bird either initiates a motif, called vocally active (light blue), or does not vocalize, called vocally inactive (purple). The (B) The power spectra in Fig. 4A are normalized and naively decomposed into PSCs (see Methods). The elements of the first principal spectral component (1^st^ PSC, blue) is non-zero across all frequencies, likely due to the power law in LFP PSDs. The 2^nd^ PSC, golden-yellow, peaks between 0-10 Hz. The 3^rd^ PSC, brown, peaks between 10-30 Hz, but has variations across birds that extend into 50Hz. As the PSCs are all eigenvectors their signs do not matter when interpreting them. This structure is largely consistent across channels and across days. Note the other PSCs are not shown. (C) Projection of both the vocally active and vocally inactive trials onto the first three PSCs for the same channels in (A) and (B). The color coding is the same as (A).

### Syllable onset is phase aligned to underlying LFP rhythms

We next calculated the Inter-Trial Phase Coherence (ITPC) of the spectrum (methods) from 2-200 Hz to determine which frequencies had stable song aligned structured changes in phase. A defining characteristic of zebra finch song is its stereotyped motif. Leveraging this stereotypy, we calculated the ITPC aligned to the first motif of all bouts recorded in a single day (n>28). Any LFP phase coherence occurring prior to song production can be attributed to the preparation of the subsequent vocalization. In two of the three birds, the first motif was almost always preceded by an introductory note. As each frequency and time sample are separate results with an attributed p-value they are all visualized in terms of their Rayleigh Z statistic (see methods) to allow for comparisons. Fig 5A shows long lasting phase structure that precedes the onset of the bout by up to 100 ms in frequencies lower than 50 Hz. This structure continues during the course of the bout and terminates soon after the end of the bout Fig 5B. Similar structures in the ITPC are observed across channels and across bird subjects [S6 Fig]. The phase coherence is highly sensitive to jitter in the aligned behaviors as shown in the more variable birds which motif was less deterministic in structure (S6 Fig).

**Fig 5:**
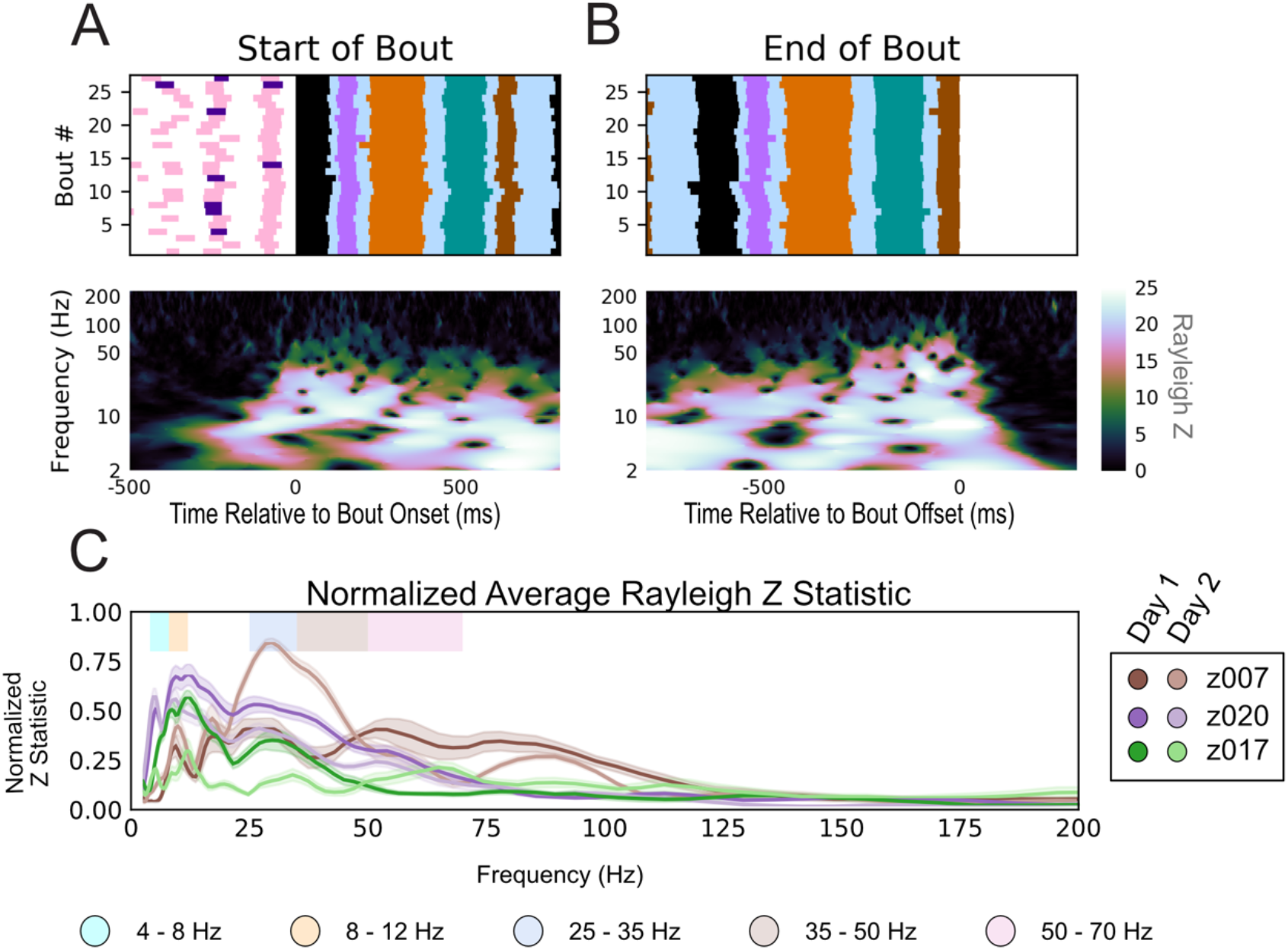
LFP Inter-trial phase coherence during production of learned sequences. (A) ITPC of LFP aligned to the start of the first motif of the bout for bird z007 (bottom) across multiple renditions of similar bouts (n=27, top) aligned to the start of the first motif in the bout. (B) the same as (a) aligned to the end of the last motif of the bout. No dynamic-time warping was used. To ensure the start and end of the bout are unique time periods, only bouts with more than one motif in duration were used. Behaviorally inconsistent bouts were excluded for clarity of visualization; including these bouts does not alter the result. (p<0.006 for all Rayleigh Z > 5). (C) Normalized sustained Z-statistic of the ITPC for samples preceding the labeled start of the bout. The steps for calculating this metric are detailed in the methods. Data reflect all bouts from both high yield days for three birds.

To determine the frequencies where phase coherence is most consistent across individual subjects, we computed the normalized sustained Z Statistic (methods) for the two high yield days for all subjects (Fig 5C). The frequency ranges that contained the peaks in the normalized sustained Z statistic were largely consistent, although the magnitude of the peaks varied. The frequencies that contained sustained high ITPC values include the 25-35 Hz range, as previously established [31,33], and several other frequencies: 4-8 Hz, 8-12 Hz, 35-50 Hz, and 50-70. These oscillations fall within well documented frequency ranges in mammalian literature namely theta (approximately 4-8 Hz), alpha (approximately 8-12 Hz), and low gamma (approximately 30-70 Hz) [54]. The lower frequencies exhibited longer periods of phase coherence through time than the higher frequencies. These periods occurred over several cycles and would fall out of alignment faster than a full cycle once song terminated (Fig 5B and S6 Fig). Strong phase coherence was observed throughout the production of the motif without the need for dynamic time warping to force motifs into the exact same time scale (Fig 5 and S6 Fig).

To better understand the dynamics underlying HVC population activity, we examined the song-aligned LFP phase coherence in greater detail. One hypothesis, given the highly stereotyped spectro-temporal structure of the motif including the brief gaps between each syllable, is that both the vocal output and the underlying HVC dynamics are largely deterministic. That is, the observed coherence may reflect an initial, single alignment at the start of the motif, that endures only by virtue of the very low song-to-song variability. Alternatively, it could be that HVC population activity reflects a more tightly controlled timescale, in which the observed oscillations are aligned to shorter timescale vocal events such as syllables. To determine whether alignment is tied to the motifs (hypothesis one), or unique to each syllable (hypothesis two), a smaller scale time alignment was used. We first verified that phase locking was found when localized to each vocalization. Syllable specific polar phase analysis, shown in Fig 6A, shows a strong phase preference to the onset of each syllable in all of the previously determined frequency bands. These phases were unique to each frequency band for each syllable. Similar results were seen for all subjects across all days (S7–S10 Fig). We also found the same phase preference to vocalization onset for the introductory note despite its huge variability in duration and structure beyond a single utterance. Fig 6B and 6C demonstrate that this stereotyped phase structure occurs prior to and during each vocalization and is statistically unlikely to have occurred by chance. To directly test the two hypotheses, we compared the ITPC centered on each syllable after the first to the ITPC centered on their stereotyped time within the motif (Fig 7). Centering on the labeled syllable start time yield significantly stronger ITPC compared to centering on the stereotyped, or averaged, start time for each syllable within the motif (S7–S10 Fig). Thus hypothesis two is the more likely source of this phase locking characteristic.

**Fig 6:**
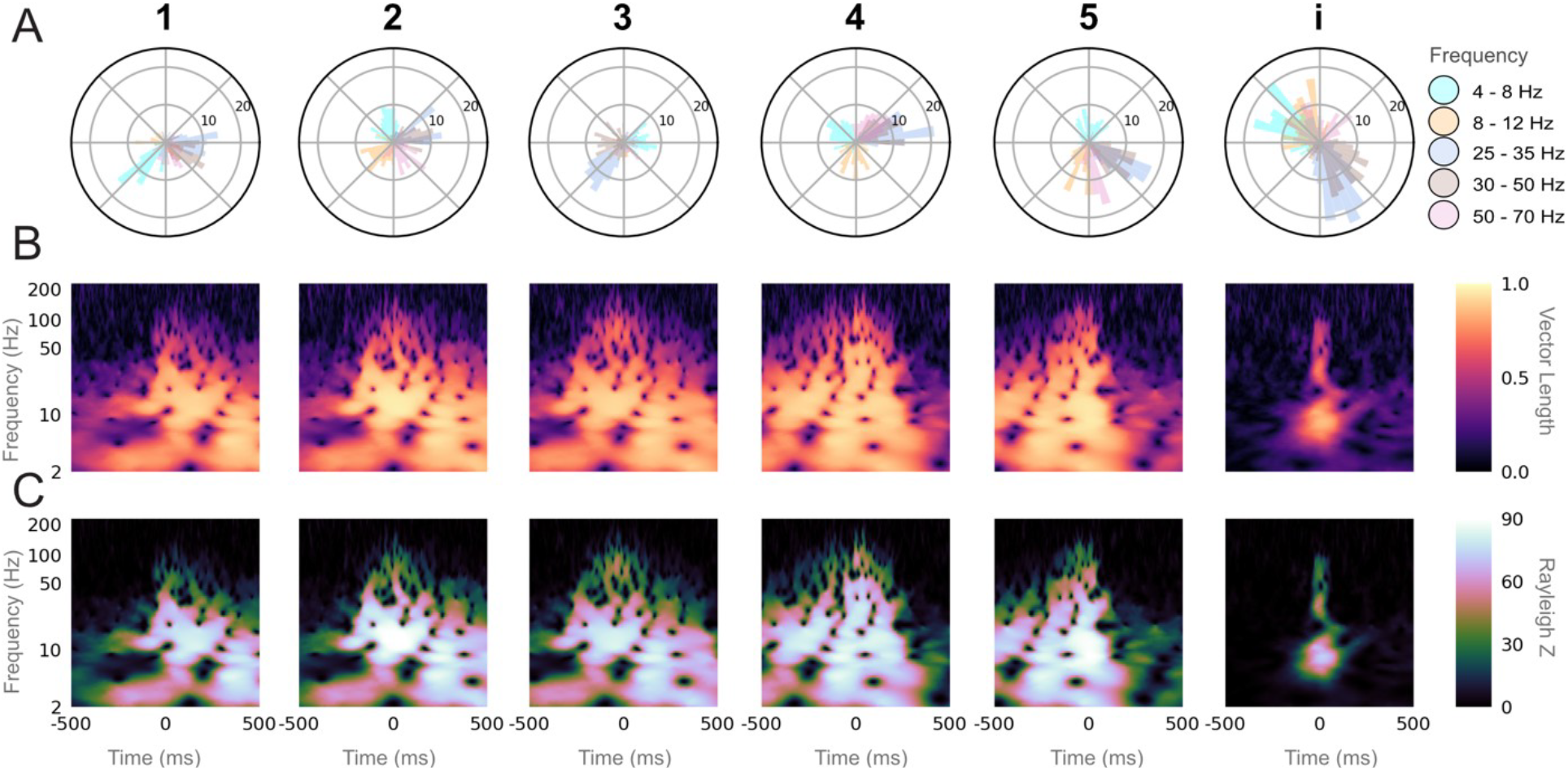
Syllable specific inter-trial phase coherence reveals phase preference to syllable onset. (A) Polar histogram of the phase for each LFP frequency band at the labeled start of all instances of a given syllable or the introductory note over the course of one day (Day 2), for one bird z007. The number of instances (n=100) are equal for all syllables and the introductory note, and is set by the syllable class with the fewest renditions. (B) ITPC resultant vector length, indicating the level of phase consistency across trials, for each frequency over time relative to the labeled start of each syllable or introductory note (0 ms) over the same instances as in (A). (C) Rayleigh Z-statistic of the ITPC over the same time and frequencies as (B).

**Fig 7:**
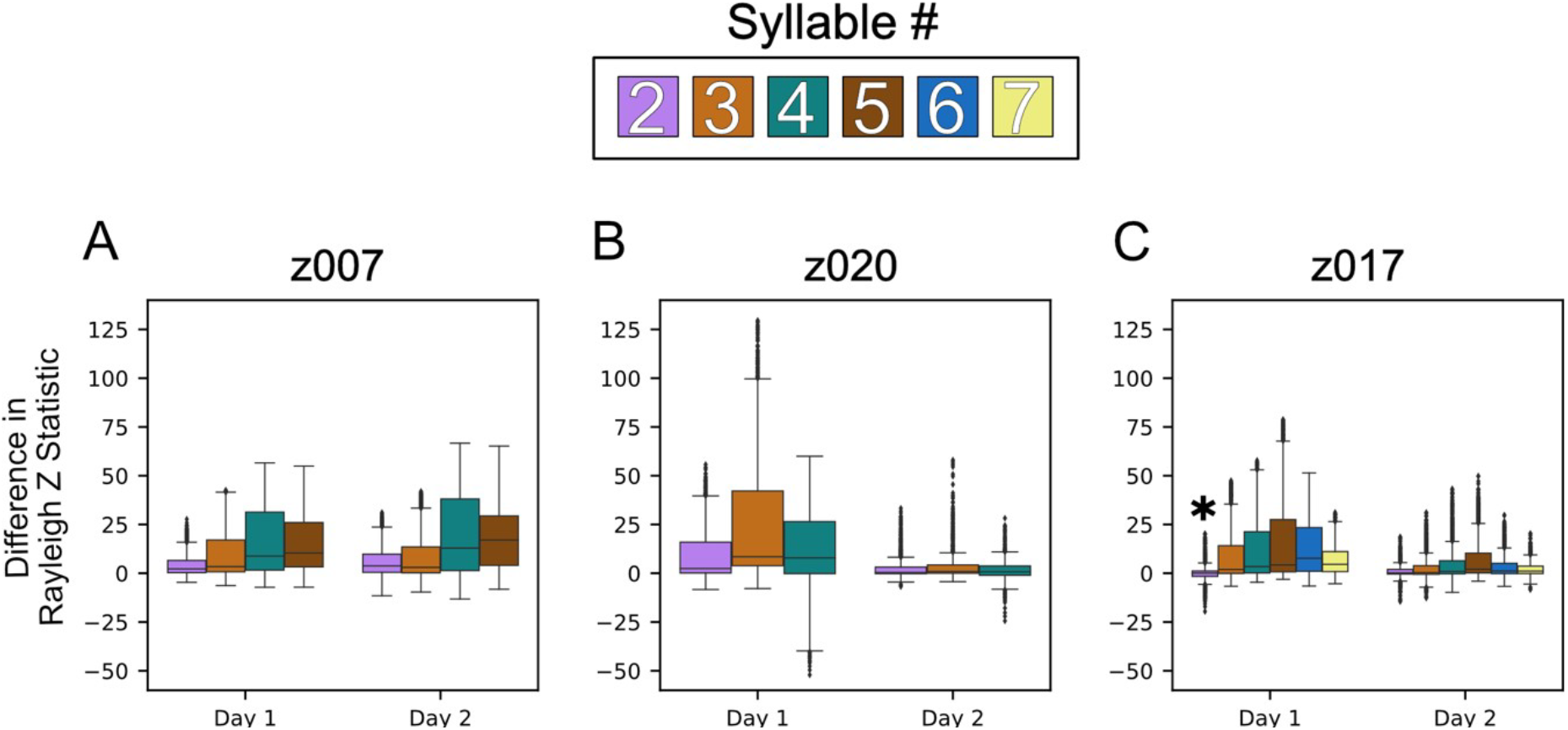
Phase preference to syllable phase preference is reflective to constitutive syllables rather than the larger motif structure. Box plots of the differences in Rayleigh Statistic between the syllable aligned ITPC and the ITPC aligned to that same syllables’ stereotyped onset time within the motif. Positive differences indicate greater phase consistency at the syllable aligned time versus that of the stereotyped time across LFP frequencies (4-8 Hz, 8-12 Hz, 25-35 Hz, 35-50 Hz, and 50-70). This difference was determined by first concatenating the Z Statistic for each channel centered at the designated time point to get a vector of all partially overlapping frequencies for all channels. The Vector of stereotyped alignment was then subtracted from the labeled onset alignment to get the difference for each frequency on every channel. This was repeated for all syllable, excluding the first, with each syllable represented with a specific color as indicated. (A) Shows the results for the two high yield days for z007, (B) is the results for z020, and (C) is the results for z017. All instances of each syllable that were preceded by the first syllable were used. To determine statistical significance the one sided Wilcoxon signed-rank test was used with each frequency and channel pair. * denotes that the comparison was **not** statistically significant when using the using the Benjamini-Hochberg False Discovery Rate. All other results p<0.05 and q<0.05.

### LFP features encode intended syllable identity

With potential LFP bands found through the decomposition of both phase and power we next asked whether their spectral features were correlated to vocalization identity. If so, then these features could be used to classify vocalization identity from neural activity alone. As the dominating characteristic of these features were there consistent structure, a promising approach was to create LFP templates that could be correlated with representative time traces. It was unclear however, what the ideal time range and latency relative to song onset for this information might be. To examine this, we conducted a hyper-parameter search using a Linear Discriminant Analysis (LDA) model using Singular Value Decomposition (SVD) (see Methods). The classifier was trained on varying combinations of template duration in samples (bin width) and the number of milliseconds prior to event onset from which the template was taken (offset). For most frequencies found, multiple combinations of offset and binwidth were able to correctly classify between song syllables, silent (non-vocal) periods, and —for the subject that performed them— introductory notes (z020 & z007) significantly above binomial chance (Fig 8 and S13 Fig). As this classifier had to learn to distinguish from both vocal and non-vocal events; they are collectively referred to as classes (see Methods). The parameter search for each frequency was done separately and found increasing performance with decreasing offset. S8 and S9 Tables summarize the parameters in the search yielding the highest classification accuracy. The information for vocal classification from lower frequencies were stable over longer time spans, i.e. larger bin widths, and may be more useful for predicting song (bout) onset. Nearly all frequencies we examined were useful for classifying prior neural activity as it relates to the song it will produce downstream; however it is not immediately clear what component of these oscillations, i.e. phase or amplitude, carries information regarding song identity.

**Fig 8:**
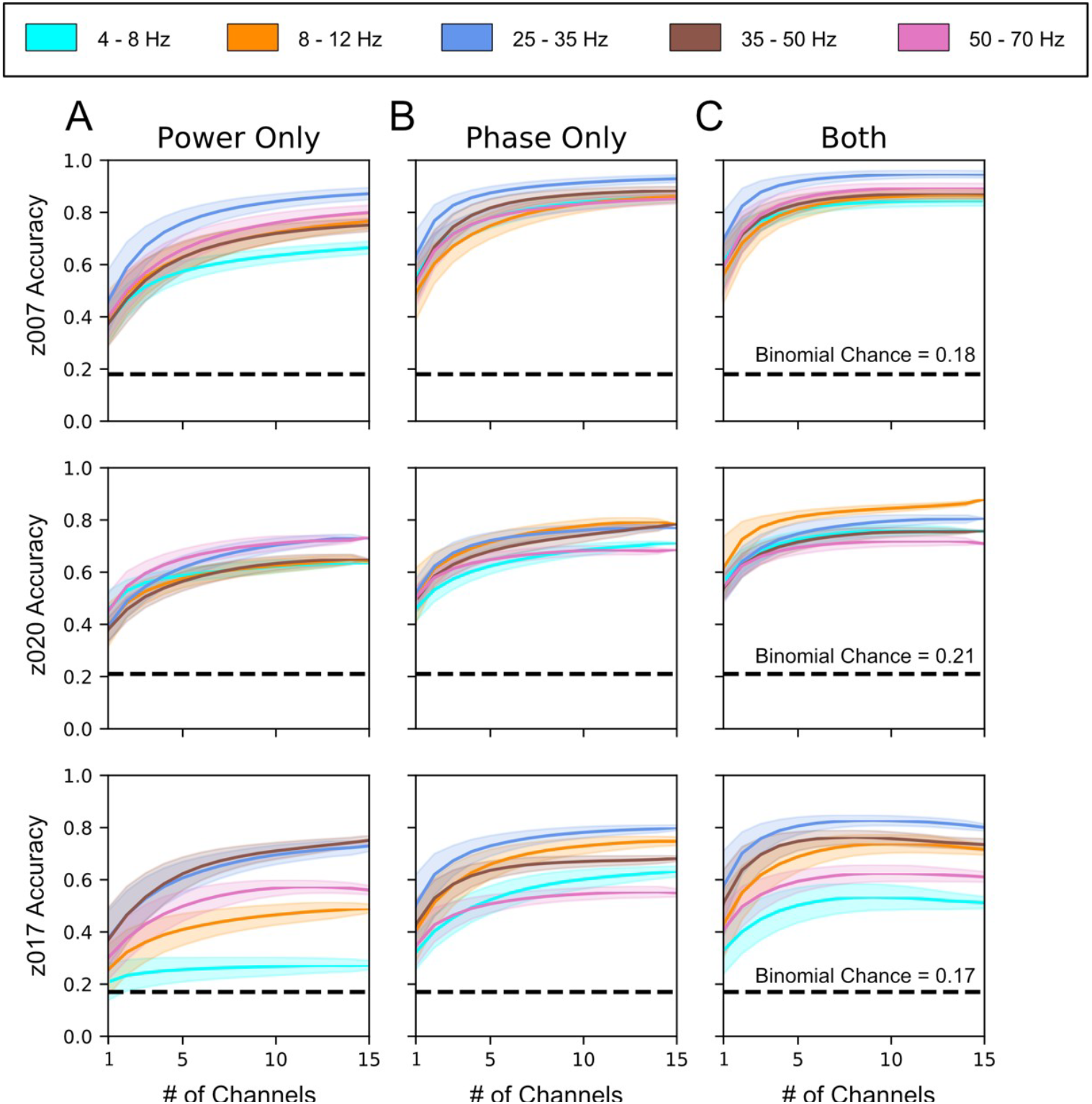
LFP phase and power provide independent and additive information for vocalization classification. Channel adding curves calculated by repeatedly training classifiers with an increasing number of randomly selected channels (see Methods). (A-C) Channel adding curves showing classifier performances with either (A) all LFP phase information removed, (B) all LFP power information removed, or (C) with both phase and power used as independent features. Each row corresponds to data from the highest yield day for each bird. These vocal and non-vocal classification events are collectively referred to as classes (see Methods). z007 n=98 for each of 7 classes (1, 2, 3, 4, 5, i (introductory note), Silence), z020 n=91 for each of 6 classes (1, 2, 3, 4, i (introductory note), Silence), and for z017 n=52* for each 8 classes (1, 2, 3, 4, 5, 6, 7, Silence). The results up to 15 channels are shown to allow for direct comparison across subjects. Dark lines show the mean for each vocalization class, shaded areas give the standard deviation over the bootstrapped analysis using n=5,000 repetitions across 5 cross-validation folds. The p-value for all of the Binomial chances calculated for each bird was 0.05. *The number of instances for each class was limited by Syllable 7, which is a intra-motif note.

To determine which component (phase or amplitude) of each frequency band provided information about the vocalization identity, we used Hilbert transforms to selectively remove either phase or power, while retaining only the contributions of power or phase information, respectively. Information provided by a given component was inferred from classifier accuracy. As shown in Fig 8A-C both phase and amplitude had accuracies above binomial chance for all frequencies and from this we inferred that they each carried song related information. In general, however, phase had higher classification accuracy than power for the frequency bands below 50 Hz. This difference in accuracy between phase and power was greater for lower frequencies (below 30 Hz). The highest frequency band, 50-70 Hz, had marginally higher performance for power only than phase only. We also asked if the number of channels recorded were necessary for decoding song identity by running a channel adding analysis for each frequency band (see Methods). This analysis also shows at what point having additional channels on the recording electrode gives diminishing improvements to the classifier performance. As shown in Fig 8A the LDA classifier performed well above binomial chance with only 5-10 channels of neural activity when classifying between song syllables, silence and introductory notes.

### Causal LFP Features Predict Syllable Onsets

The ability to accurately classify specific syllables using the features of the LFP, may be possible because a unique LFP structure associated with specific syllables or because both tend to co-occur at specific times within the motif. To dissociate these two possibilities, we examined whether the features optimized for determining syllable identity (S8 and S9 Table) could also predict specific onset times of each separate vocalization within the full motif. To do this, we implemented a naive pattern-matching approach (Methods). Because of the well-documented and well-studied stereotyped structure of zebra finch’s song, the mean error between the stereotyped start time of each syllable—relative to its first syllable—and the actual time the syllable occurred was used as a benchmark to test the performance of a predictor that used the LFP features as described before. An example of one motif and its corresponding confidences is shown in Fig 9A. Across all syllables for all birds the predictor that used all of the frequencies performed better than using the deterministic onset of the syllable only, Fig 9B. Similar performance was found for the 2 additional syllables for z017 as well (S15 Fig). These results mostly hold true across all days analyzed and were consistently found to be statistically significant (p<0.05) using the Wilcoxon signed rank test. This paired test directly compared the error between either the neural predictor or the behaviorally based predictor to the true onset time to determine whether the neural based predictor produced less prediction error. There were only three cases in which the LFP predictor’s performance failed to meet significance when using the Benjamini-Hochberg False Discovery Rate; they were syllable 2 & 4 for z020’s 2nd high yield day and syllable 2 for z017’s 2nd high yield day (S15 Fig). We next asked which frequency, if any, is best at predicting the onset of the syllables. As shown in Fig 9C, no single frequency performs best for predicting onset times across all frequencies and all subjects. Although the higher frequencies tend to do better and have lower variance than the lower frequencies, using all of the frequencies yields better performance than any one frequency across all subjects and sessions. There were poor prediction performances for 4-8 Hz, 8-12 and the 50-70 Hz for some syllables for certain birds. Further, results from 4-8 Hz were more likely to be statistically insignificant and performed worse than the stereotyped reference for most syllables.

**Fig 9:**
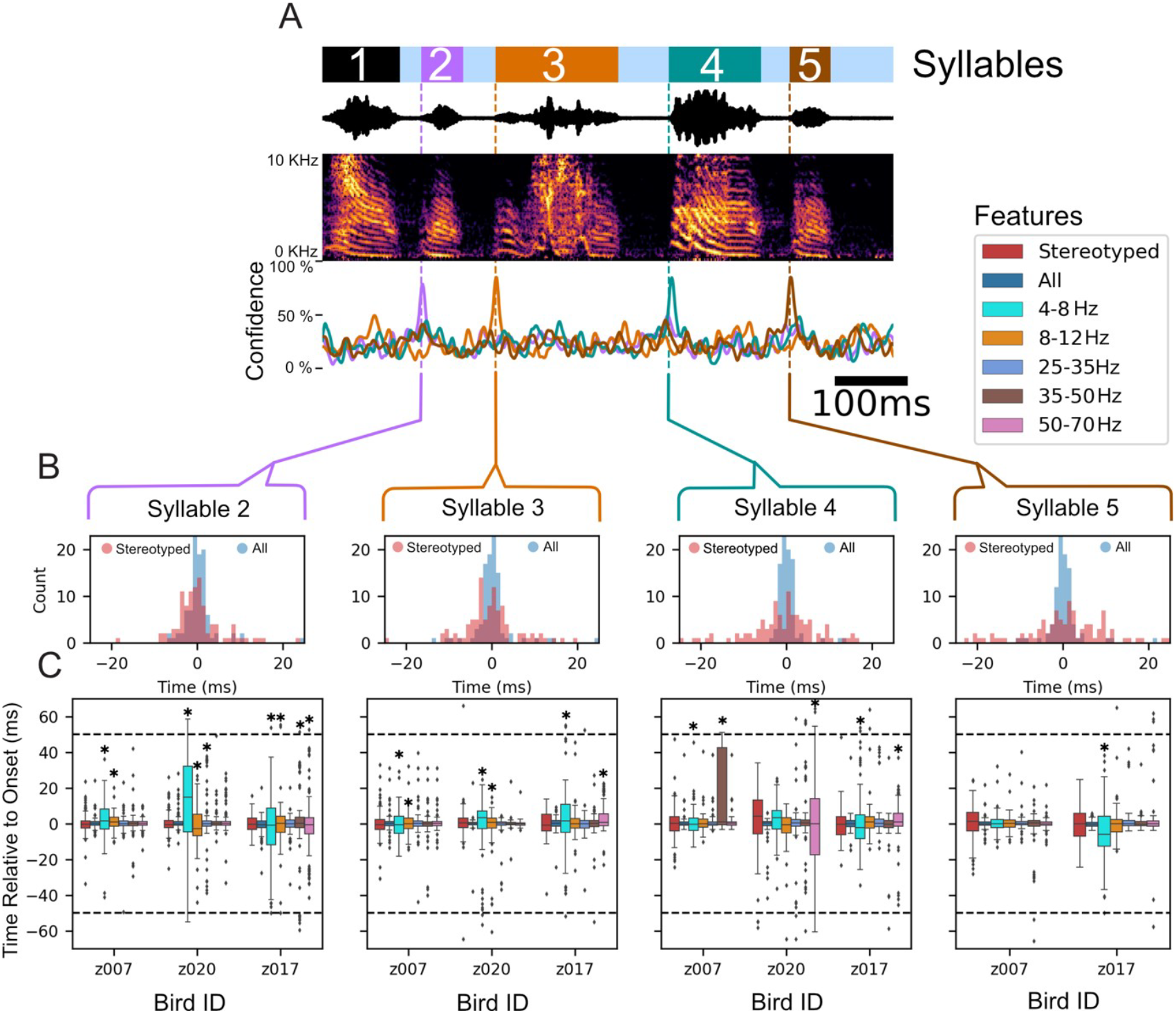
Onset detection using LFP features. (A) Example motif from the highest yield day for subject z007. Annotated behavior (top) using the same color scheme as in Fig 2, sound pressure waveform and the corresponding time-aligned spectrogram (middle), and the time-varying naïve confidence of the onset prediction (bottom) for each syllable in this example motif. Confidence signal traces are the same color as the syllable they are meant to predict. (B) Histogram of onset prediction times for each syllable relative to its annotated start time. The annotated start times are relative to the start of the first syllable of the motif the syllable occurred in. The histogram compares two approaches. The first, in pink, uses only behavioral information (the stereotyped onset relative to the start of the first syllable). The second, in blue, uses all the neural features to predict the start of the syllable. (C) Boxplot of onset prediction times relative to the labeled onset time for each bird during its highest yield day. The order of each feature used is the same, going left to right: first is the stereotyped onset time using only the deterministic behavior, next is the results using all of the neural features, then each frequency band only in order from least to greatest (4-8 Hz, 8-12 Hz, 25-35 Hz, 35-50 Hz, and 50-70 Hz). The time window that the neural based predictor must make a prediction within is represented by the dotted black line represents (see Methods). Statistical significance was calculated using the one sided Wilcoxon signed-rank test, and * denotes results that are **not** statistically significant when using the Benjamini-Hochberg False Discovery Rate. All other results p<0.05 and q<0.05.

## Discussion

We demonstrate that the local field potentials in putative zebra finch HVC carry significant information tied to vocal production. More specifically, the time-varying structure of this meso-scale signal, captured by both the power and phase of various LFP frequency bands, correlates with behaviorally-relevant timepoints during this complex, learned motor behavior. Our results extend previous studies in zebra finch HVC [33], by showing that LFP features can be used to detect, using simple methods, both when and what syllable the bird will sing. Prior to this previous study and our work, little was known about the relationship between LFP features and vocal production in birds. This limited the generalizability of the otherwise powerful birdsong model to other systems, including non-human primates and especially humans, where LFP and surface potentials are broadly used to measure the dynamics of cortical motor control [12,47,50,55]. In addition, we note that the bandwidth and features of the LFP signals investigated in this paper share similarities with LFP features tied to motor control in humans, non-human primates, and mammals. For example, the power spectrum components most closely tied to song in finches (Fig 4 and S5 fig), match those documented in the human motor cortex during finger flexion [50]. We suggest that LFP recordings can serve as useful targets to further our understanding of songbird neurophysiology, and to more closely connect the powerful birdsong model to motor control studies in mammals, non-human primates and humans.

A striking feature of the amplitude changes in LFP during song is the increase in higher frequencies that are within the band often referred to as “High Gamma”. Although changes in this frequency band in Nif have been previously reported [56], they were not found to be premotor related. We have shown the first preliminary evidence, to our knowledge, of premotor activity related amplitude changes in high gamma (S2 Fig). This is significant because this band is often the feature of choice for many state of the art speech BCIs in humans [12,24]. Notably these changes empirically appeared to correspond introductory notes in addition to song syllable. Future work should evaluate if these amplitude changes also occur during calls which are considerably less deterministic and syntactically complex than the zebra finch motif.

A consistent feature observed in the LFP of putative HVC in songbirds during singing is tight phase locking to vocal behavior. In addressing the question of what potentially gives rise to the observed song-aligned phase locking; we proposed two hypotheses. Hypothesis one where the phase locking is related to the larger motif sequence and hypothesis two which attributes the phase locking to each unique vocal units (syllables). In simpler terms is the phase locking a continuous process aligned to the start of the motif (Hypothesis one) or discrete process that aligns to each syllable (Hypothesis two). When directly testing the two hypotheses the second proved more statistically likely. While there is noticeable non-deterministic jitter in the brief gaps of silence between syllables that make up the motif, this may not be enough to completely disprove hypothesis one. However there are other notes that occur non-deterministically between motifs, referred to as intra-motif notes or “connectors” [57] and introductory notes for whose timing before the motif is variable. Looking at the phase polar plots of individual syllables would either show high selectivity of phase preference for all unique vocalizations if hypothesis two is correct or low levels of phase locking for motif syllables and a random distribution about the polar axis (null) for the intra-motif notes and introductory notes if hypothesis one is correct. As shown in Fig 6 and Fig 7, each syllable has phase preference that is stable across every specific vocalization type including intra-motif notes (z017 & z020) (S8–S10 Fig) and introductory notes (z020 & z007) (Fig 6 and S7 Fig). This result indicates that precise phase preference is behaviorally relevant to both learned and non-learned vocalizations. Thus, it can be thought that the long template phase traces in lower frequencies that precede and persist during song production are a byproduct of the zebra finches’ stereotyped song structure. This structure is composed of encoding at a smaller time scale centered on each individual syllable during song production. Similar phase preference to motor onset has been found in both human and non-human motor cortex [58].

When showing the results of the normalized sustained Z statistic there was a difference in in magnitude of peaks both between and within birds. This is due to there being significant variation in both these dimensions. This variation likely has many sources, including electrode placement and varying signal to noise ratios across days. Within days the peaks in the Z-statistic denote consistency in phase locking. Although the magnitude of these peaks varied, the frequency bands that contained them were empirically similar to those described in literature. These result, existing literature, and empirically inspecting narrowband, as show in S3 and S4 Fig, informed our narrowband feature selection.

Previous work in Zebra Finch from Markowitz et al showed that spiking of both excitatory and inhibitory neurons were locked to the phase of the 25-35 Hz band [31,33]. Admittedly this previously identified phase locking combined with the behavior-locked manner of spiking activity to song behavior point to some of the phase locking we observed. However, our work shows that other frequency bands beside the 25-35 Hz band are phase locked to the behavior. More importantly we find that these frequencies, and the 25-35 Hz band, are also locked to other behaviors besides just the syllables of the motif, namely intra-motif notes and introductory notes. This potentially opens the door to additional vocalizations in the zebra finches repertoire to be used to study motor-vocal control. These behaviors are significantly less deterministic than the syllables of the motif. Admittedly, the stereotyped structure of the Zebra Finch’s song differs from the more complex structure of human speech. Broadening the vocal behaviors that can be used in this type of behavioral decoding analysis can help mitigate this weakness in the zebra finch model. In addition, the methods and insights learned from them can be directly applied to songbirds with more complex song structures such as starlings, canaries, and bengalese finches. Collectively these songbird models give the opportunity to investigate vocal-motor encoding at varying levels of behavioral complexity.

Naturalistic motor behavior is intrinsically difficult to study due to its high degree of variability and limits on brain coverage in chronic experiments. Zebra finch are a desirable model species because they mitigate the first of these difficulties. Their song allows for repeated segments of the same sequence to be produced with two almost opposing characteristics: near perfect precision of each utterance and the natural irregularities that exist in all of nature. This provides a dataset that has both high repetitions of each syllable class and a non-deterministic structure beyond the motif level that facilitates detailed analyses. This model is a unique testbed for approaches to predict the onset of vocalizations in other animals, particularly humans. The simple onset prediction results where each syllable after the first were predicted using causal neural features, described in Fig 9, were done using little-to-no parameter optimization and take into account no information regarding the structure of the song. Better performance could be achieved with more elegant approaches with minimal increases in computational complexity. Suggesting that these features could be leveraged to predict song onset with low latency. At present the state of the art for Human speech BCI’s uses non-causal neural signals to synthesize sounds similar to the intended speech [12,24,59]. While promising, the non-causal nature of these algorithms introduces significant latency [59] and ultimately limits the quality of interaction with such a prosthesis. Even further, the limits on how long researchers have access to neural signals impedes the speed at which these offline techniques can be translated in real-time in a clinical setting. This is where the songbird model as a whole can contribute to the BCI field; as an animal model to proof-of-concept systems in which closed-loop interaction with prosthesis designs can be rigorously studied. Systems derived from these proof-of-concepts will be highly valued when translated to the human clinical setting. These features and approach provide a starting point for further analysis to look to zebra finch and other songbirds as more than a model for vocal learning and sequence generation, but as a model of vocal prediction and BCI development.

## Methods

We implanted male zebra finch with laminar electrodes and simultaneously recorded their vocal behavior and neural activity (Fig 1). Through a series of steps, explained in detail in the Annotation and Alignment of Behavioral Data section, we found segments of the recording that contained songs and hand annotated them using Praat (Fig 2). These behavioral labels were used to analyze the neural activity in relation to specific classes of behavior to determine what frequencies, if any, correlated to the behavior (Fig 3–6). Finally, the frequencies discovered were used to classify and predict the onsets of the behaviors to clarify their relationship to the vocalizations (Fig 8 & 9).

### Subjects

All procedures were approved by the Institutional Animal Care and Use Committee of the University of California (protocol number S15027). Electrophysiology data were collected from three adult male zebra finches. Birds were individually housed for the entire duration of the experiment and kept on a 14-h light-dark cycle (Fig 1A). Each day had one session that lasted multiple hours and were unstructured as they depended on the subject’s willingness to sing (Fig 2A). All available days were analyzed, however the two highest motif yielding days, hereafter referred to as *high yield days*, were reported and used for statistical analysis. The full duration of chronic recordings ranged from 5 to 19 days. The birds were not used in any other experiments.

### Electrophysiology and audio recording

Electrophysiological recordings were gathered from 3 subjects, in which two were implanted with 16-channel probes and one with a 32-channel probe (Fig 1A). We used 4-shank, 16/32 site Si-Probes (Neuronexus), which were PEDOT-coated in-house (https://www.protocols.io/view/EDOT-PSS-c2syed). The probes were mounted on a custom designed printable microdrive [60] and implanted targeting nucleus HVC (Fig 1A & B). Audio (vocalizations) was recorded through a microphone (Earthworks M30) connected to a preamplifier (ART Tube MP) and registered temporally to ongoing neural activity (Fig 1)]. Extracellular voltage waveforms and pre-amplified audio were amplified and digitized at 30 kHz using an intan RHD2000 acquisition system, Open Ephys and custom software (Fig 1D) [61].

### Electrode implantation procedure

Animals were anesthetized with a gaseous mixture of Isoflurane/oxygen (1-2.5%, 0.7 lpm) and placed in a stereotaxic frame. Analgesia was provided by means of a 2mg/kg dose of carprofen (Rimadyl) administered I.M. The scalp was partially removed and the upper layer of the skull over the y-sinus was uncovered. The probe was attached to the shaft of a microdrive of our design (https://github.com/singingfinch/bernardo/tree/master/hardware/printable_microdrive) which was printed in-house using a b9 Creator printer and the BR-9 resin. A craniotomy site was open 2400 μm medial to the y-sinus (right/left hemispheres). The dura was removed and the electrode array was lowered to a 300-500μm depth. The opening was then covered with artificial dura (DOWSIL 3-4680 Silicone Gel Kit) and the microdrive was cemented to the skull using dental cement (C&B Metabond). A reference wire was made with a 0.5 mm segment of platinum-iridium wire (0.002”) soldered to a silver wire lead and inserted between the dura and the skull through a craniotomy roughly 3mm medial (contralateral to the hemisphere where the electrode was inserted) and 1.5 mm anterior to the y-sinus. The reference electrode was cemented to the skull and the silver lead was soldered to the ref and gnd leads of the neuronexus probe. The craniotomy, the electrode, and the Microdrive were then covered with a chamber designed and 3D printed in house, which was cemented to the skull. The skin incision was sutured and adhered to the chamber with adhesive. The mass of the probe, microdrive and protective chamber were measured to be 1.2-1.4g. Upon returning to a single-housing cage, a weight reliever mechanism was attached using the end of a thin nylon wire that was attached to an ad-hoc pin in the chamber; the other end routed through a set of pulleys and attached to a counterweight mass of ~1g [61].

### Analysis of electrophysiology data

Extracellular voltage traces were multiplexed and digitized at 30kHz on the headstage, and stored for offline analysis. **--offline--** They were then low-passed filtered at 400 Hz using a Hamming finite impulse response (FIR) filter and downsampled to 1 kHz. The group delay introduced by the filter is compensated by introducing a temporal shift to the filter output [62].

### Annotation and alignment of behavioral data

The song of an adult male zebra finch can be partitioned and labeled in multiple ways. However, the most fundamental characteristic of their song that is agreed upon is their stereotyped motif. The motif is a consistent sequence of sounds interleaved with silent periods that is learned from a tutor while they are young [10,29,30]. Song motifs are also grouped into larger structures called bouts, which consist of multiple repetitions of motifs [10,29,57]. Depending on the definition used, bouts can include a short repeated vocalization that typically precedes the first motif of the bout. For the purpose of the analysis done, the start of the bout is explicitly defined to be the start of the first motif in that bout. This is due to the focus of understanding the correlation between LFP and learned motor sequences. Beyond the syllables of the motif male zebra finch may have an additional syllable or sequence of syllables they will optionally insert between successive repetitions of motifs. This is called a “connector” [57] or intra-motif note. In our recordings both z017 and z020 had intra-motif notes. Because z007 did not have any intra-motif notes, and therefore a more stereotyped song, it will be used for all of the empirical figures shown.

A custom template matching algorithm written in Python was used to find potential instances of vocal activity (using an exemplar motif as the template), which were then curated manually to rule out false positives [61]. The curated motif start times were grouped into larger time segments that ensured that the gap between each motif was no greater than 20 seconds (Fig 2A and 2B). These chunks of time are subsequently referred to as vocally active periods (VAP). These VAPs were then manually labeled using the Praat speech analysis software [63]. Vocal events were segmented by hand (Fig 2C) and labeled based on identity. There is a unique label for each individual syllable and introductory note (if the bird demonstrated one). All other male vocalizations were grouped together and labeled as a call. Calls are short, simple vocalizations that are produced by both sexes which mostly do not have learned components [30]. There were also two labels for silence in these VAPs: one for the gaps between song syllables, and another for silence not during song in the VAP.

To leverage these behavioral annotations for contextual analysis of their synchronously recorded neural activity, custom Python software was developed [BirdSongToolbox]. This software added additional labels to the hand labels based on their sequence and context within the larger bout structure; e.g. first or last motif in bout. Labeled vocalization segmentation times were adjusted to the proper sampling rate for the downsampled neural data using this same software. Finally, the software used these additional contextual labels to select particular event times to align both vocal and neural data that fit certain criteria matching the particular vocalization of interest.

### Time series power spectrum

Spectrograms were calculated by first filtering the data with 100 partially-overlapping narrow frequency bands that were equal width in log-space. These filtered time series were Hilbert transformed to extract the analytic amplitude at each time point. Each narrowband amplitude time series was then normalized by the mean of that frequency band over the entire duration of its VAP.

### Principal spectral decomposition

Following work done in Human ECoG work, a principal component method was applied to find consistent changes in LFP spectrum related to motor-vocal behavior. This method required the calculation of the power spectral density (PSD) for windows of time during either vocal activity or inactivity. Although the behavioral changes in birdsong are on a smaller timescale (20-100 ms) than human limb movements, to keep methods consistent between studies the same window size of 1 second was used for the trial length, *τ*_*q*_. PSDs of vocally active trials centered on song activity, which included introductory notes and calls, were analyzed alongside trials of neural activity centered within larger periods of vocal inactivity longer than 2 seconds in duration. All PSDs were calculated using the multitaper method with the time_frequency.psd_array_multitaper function from the MNE-Python software package [62]. The frequency range was set from 0 to 200 Hz with 14 orthogonal slepian tapers. Each PSD, *P*(*f*, *τ*_*q*_), was individually normalized using two steps: each spectral sample was elementwise divided by the average across the ensemble, at each frequency, and then the log was taken.

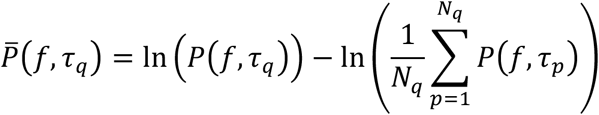

The label *q* refers to the times centered within periods of high vocal activity and vocal inactivity (silence). The number of instances per class were balanced to be equal and their combined total number of PSDs is denoted by *N*_*q*_. The covariance matrix 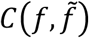 between frequencies of these normalized PSDs were calculated: 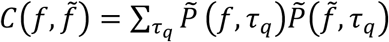. The eigenvalues, *λ*_*k*_, and eigenvectors, 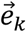, of this matrix elucidate common features during song production. These eigenvectors, 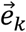, are referred to as “Principal Spectral Components” (PSCs) [50].

#### Cosine Similarity

Cosine similarity is a measure of similarity between two non-zero vectors of an inner product space. For the purpose of comparing the principal spectral components it evaluates the orientation, not magnitude, of two vectors in relation to one another. To calculate it you take the inner product of two unit vectors, or two vectors whom have both been normalized to have a length of 1.

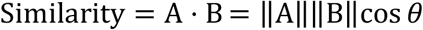

To calculate the cosine similarity for the PSCs we must take the dot product between two unit vectors. As the PSCs are all eigenvectors calculated using PCA they are already unit vectors and their sign can be flipped without altering their information. A template of each PSC is calculated by taking the mean of the PSCs across all good channels after all of their signs have been aligned. This template PSC is then normalized to have a length of one by dividing it by its norm. The cosine similarity matrix is then calculated by taking the dot product of these two unit vectors. This is expressed below with the template PSC represented as T and the PSC symbolized by P from a selected channel, c.

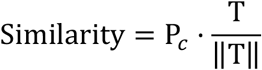

### Feature extraction

Feature extraction from the preprocessed data consisted of a series of steps that involved first Common Average Referencing and then narrow band-pass filtering. Common Average Referencing has been shown to minimize or remove the contributions of correlated noise sources such as 60 Hz noise and intermittent noise sources such as movement artifacts [64]. We used the Hilbert transform to compute the analytic signal. To extract oscillatory power the absolute value of the analytical signal was used, which effectively removed all phase related information. To extract oscillatory phase the instantaneous angle of the complex argument of the analytical signal was used, and was further smoothed using a sine function. This effectively removed all information related to power from the signal.

These signals, whether Hilbert transformed or not, were then sub-selected at times relative to the label onset for a syllable using two parameters: bin width and offset. The bin width is the number of samples used prior to the offset, and the offset is the number of samples, or milliseconds, prior to the labeled start time to a specific syllable. All combinations of these hyperparameters used offset which were prior to the start of the vocal behavior.

#### Band templates and Pearson correlation features

Using the optimized bin width and offset for a particular frequency band, a template for that band was calculated by taking the mean of the LFP traces of the training set of a specific behavior class. This template represents the average LFP activity prior to and during the production of a particular vocal behavior. This template was then used to extract features from a narrow-band LFP trace by taking the Pearson correlation of the template from a segment of neural activity of the same length in samples. This correlation value is set between −1 and 1. For the behavioral classification results, a segment of the corresponding LFP frequency band that is set by the optimized bin width and offset is used as the feature. For the onset detection analysis, the Pearson correlation for each sliding segment equal in length to the optimal bin width is used to detect behavior that corresponds to its optimal offset.

### Computation and statistical testing of phase locking across renditions

We computed the Inter-trial phase coherence (ITPC) to assess trial-to-trial synchronization of LFP activity with respect to time-locked syllables within the motif. The ITPC is a measure of the phase synchronization of neural activity in a given frequency, calculated relative to critical event times that are repeated over an experimental paradigm. To calculate the ITPC across trials, for each channel, data were first filtered in 100 partially-overlapping narrow frequency bands that were equal width in log-space. These filtered time series were then Hilbert transformed to extract the analytic phase at each time point. Then, at each time point, for each channel and frequency band, we calculated the phase-consistency across trials to estimate the ITPC. Here, ‘trials’ were defined as the start or end of a specific type of vocal event, which could be further refined by its context within the bout, e.g., first or last in bout. Once all instances of the event of interest were selected, a window of time centered on each was implemented. The ITPC for each frequency and time sample pair must be calculated independently. The equation for calculating the ITPC is described as below, for n trials, if, F_k_(f, t) is the spectral estimate of trial k at frequency f and time sample t

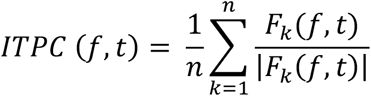

where |*x*| represents the complex norm of *x* [65]. The spectral estimate, F_k_(f, t), is the instantaneous phase of frequency f at time sample t. This was repeated for all narrow-band frequencies and time samples within the event window. The ITPC value, or resultant vector length, scales from 0 to 1, with 0 being the null hypothesis where phases are uniformly distributed about the polar axis and 1 being perfect synchrony. To determine the significance of this vector length, and to determine if it could have been randomly sampled from a uniform distribution by chance, p-values were calculated for each frequency and time sample. The mean resultant vector of the instantaneous phase across trials for a specific time sample, and its corresponding bootstrapped p-value, were calculated using the pycircstats toolbox. To enable visualization of the results for comparing ITPC values while accounting for all of the frequency and time sample combinations, the Rayleigh Z Statistic was calculated.

#### Rayleigh statistic

The ITPC over the time length of a syllable or motif is evaluated in terms of the Rayleigh Z Statistic and is defined as 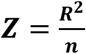 where R is the Rayleigh’s R, ***R* = *nr*** where r is the resultant vector length and n is the number of trials. P-values were estimated using the following equation **[66–69]**, 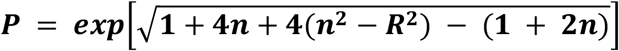.

#### Normalized sustained Rayleigh Z statistic

To determine which, if any, frequency bands could potentially be used as a feature to decode vocal behavior, a measure of how consistently significant a band stayed phase locked was created. As we don’t expect the Rayleigh Z statistic across both channels and frequencies to be the exact same, we wanted to understand the relative significance across birds and channels. The measure ranks how consistently phase locked across time a frequency was during song production overall for one recording session. The Mean Rayleigh Z-Statistic of samples over the course of one oscillation for a set frequency normalized by the maximum Z-Statistic value for the entire time period analyzed was used and will be referred to as the Normalized Sustained Z Statistic. This analysis informed the frequency bands used for the classification and onset detection analysis.

### Linear classifier

To classify behavior using the LFP features, a Linear Discriminant Analysis (LDA) model using Singular Value Decomposition (SVD) was trained using the scikit-learn toolbox [70] in Python. This classifier is tasked to correctly classify examples of all vocalizations during song, both motif syllables and intra-motif notes, in addition to introductory notes and periods of silence. As the classifier has to learn how to distinguish from both vocal and non-vocal events; they are collectively referred to as classes. All priors for each class were set equal and all classes were balanced in the datasets used for classification analyses. Detailed information regarding the number of instances in each class can be found in S3-S5 Table. No shrinkage or regularization were used, however, the SVD optimizer was used to avoid ill-conditioned covariance matrices. Results were validated using 5-fold cross-validation. Templates for feature extraction were created by taking the mean across the training set. These templates were used to extract features from both the training set and the testing set. All frequencies were trained and tested independently of one another.

#### Channel Adding Analysis

Channel adding curves were calculated using a bootstrap approach to determine how many channels were needed until additive information saturation. The channel adding curves were calculated by first training and testing a classifier with the neural features of only one channel, using the steps described previously, then repeating this analysis after adding the features of a another randomly selected channel. This repeated training and testing after adding a random channel is completed once a classifier that uses the features from all available good recording channels is evaluated. This channel adding analysis was repeated 5,000 times, changing the order in which each channel was included, with 5 fold cross validation. The order in which channels were added over the 5,000 repetitions was maintained across the folds to enable fair calculation of their validated mean. These are subsequently used to calculate the mean and standard deviation across repetitions.

### Syllable onset detection

In order to analyze the temporal relationship between LFP and song production, we used a template matching approach to determine whether LFP can predict syllable onset. Each syllable of the motif excluding the first, had every labeled instance recorded in the same day split into training and testing sets using a 5-fold stratified split (80% training set and 20% testing) (S3-S5 Table). Both the template and the stereotyped onset of the syllable were calculated from the training set. Templates are the mean of the LFP traces of the training set. The stereotyped onset is the average time the syllable occurs in the training set with respect to the first syllable of the motif it occurred in. These templates were then used to compute the Pearson correlation across time for each of the motifs that contain the syllables of the test set, maintaining the temporal relationship of the optimal offset and the bin width for its respective frequency.

The prediction confidence of a single frequency band was calculated by first thresholding the Pearson correlation values at zero, and taking the average of the resultant time series across channels. The maximum confidence within a 100 millisecond window centered on the stereotyped onset time is then used as the frequency’s prediction for that instance of the syllable. To get the results of a predictor that uses all of the frequencies the same steps as previously stated are followed; only the prediction confidence across all frequencies and channels is used. The prediction of the syllable onset using the bird’s stereotyped behavior was determined by adding the stereotyped onset time, calculated from the training set, to the actual start of the first syllable for the motif the syllable occurred in. It is important to note two things with this approach; the first is that neither approach receives information on the actual start time of the syllable in its respective motif. The second is that this 100 millisecond window is significantly larger than the natural variability of syllable onsets and gives the neural based predictor a more difficult task than the baseline stereotyped comparison. Both predictions were then normalized by taking the difference between the predicted time and the actual labeled start time of the syllable within the same motif.

Statistical significance of the results was calculated with a one-sided Wilcoxon signed-rank test using the difference between the relative predictions of the stereotyped behavioral model and the neural features models. Two null hypotheses were tested, first that there was no difference between the stereotyped behavior and the prediction using neural features, and the second being that the predictor using the neural features was closer to the actual labeled onset time. The second null hypothesis requires that the LFP based predictor must outperform the behavior based predictor. Results must pass both tests with a p-value less than .05 to be considered significant. The Benjamini-Hochberg procedure was used to control the False Discovery Rate to account for the multiple comparisons done. The procedure was implemented for each day treating the results of each individual frequency and the predictor which uses all frequencies as separate results (n=6, q=.05).

## Acknowledgements

We would like to thank the Gilja, Gentner and Voytek Labs for feedback to help refine the manuscript. We would like to especially thank Dr. Will Liberti III, Anna Mai, Abdulwahab Alasfour, Tim Sainburg, Richard Gao, and Dr. Natalie Schaworonkow for insightful comments and suggestions for improving the manuscript.

This work was supported by the U.S. NIH National Institute on Deafness and Other Communication (R01DC008358, R01DC018446, R01DC018055), NIH National Institute of General Medical Sciences (R01GM134363), National Science Foundation under grant BCS-1736028, the Kavli Institute for the Brain and Mind (IRG #2016-004), the Office of NavalResearch, Pew Latin American Fellowship in the Biomedical Sciences (E.A.), Halıcıoğlu Data Science Institute Fellowship (J.I.C.), University of California – Historically Black Colleges and Universities Initiative (D.E.B.). This material is based upon work supported by the National Science Foundation Graduate Research Fellowship under Grant No. DGE-1650112. Any opinion, findings, and conclusions or recommendations expressed in this material are those of the authors(s) and do not necessarily reflect the views of the National Science Foundation.

## Supporting information

**S1 Table:** Overall Description of Subject Data.

| Overall Description |  |  |  |  |  |
| --- | --- | --- | --- | --- | --- |
| bird_id | # of Days Recorded Total | # of uniquely Labeled Syllables in Song (Including Intra Motif Notes) | # of Intra Motif Notes in Song | Presence of Unique Introductory Note | Electrode Geometry (Neuronexus Probes) |
| z020 | 5 | 4 | 1 | Yes | A4x1-tet-3mm-150-121 |
| z007 | 9 | 5 | 0 | Yes | A4x4-tet-5mm-150-200-121 |
| z017 | 19 | 7 | 2 | No | A4x4-3mm-50-125-177 |

**S2 Table:**
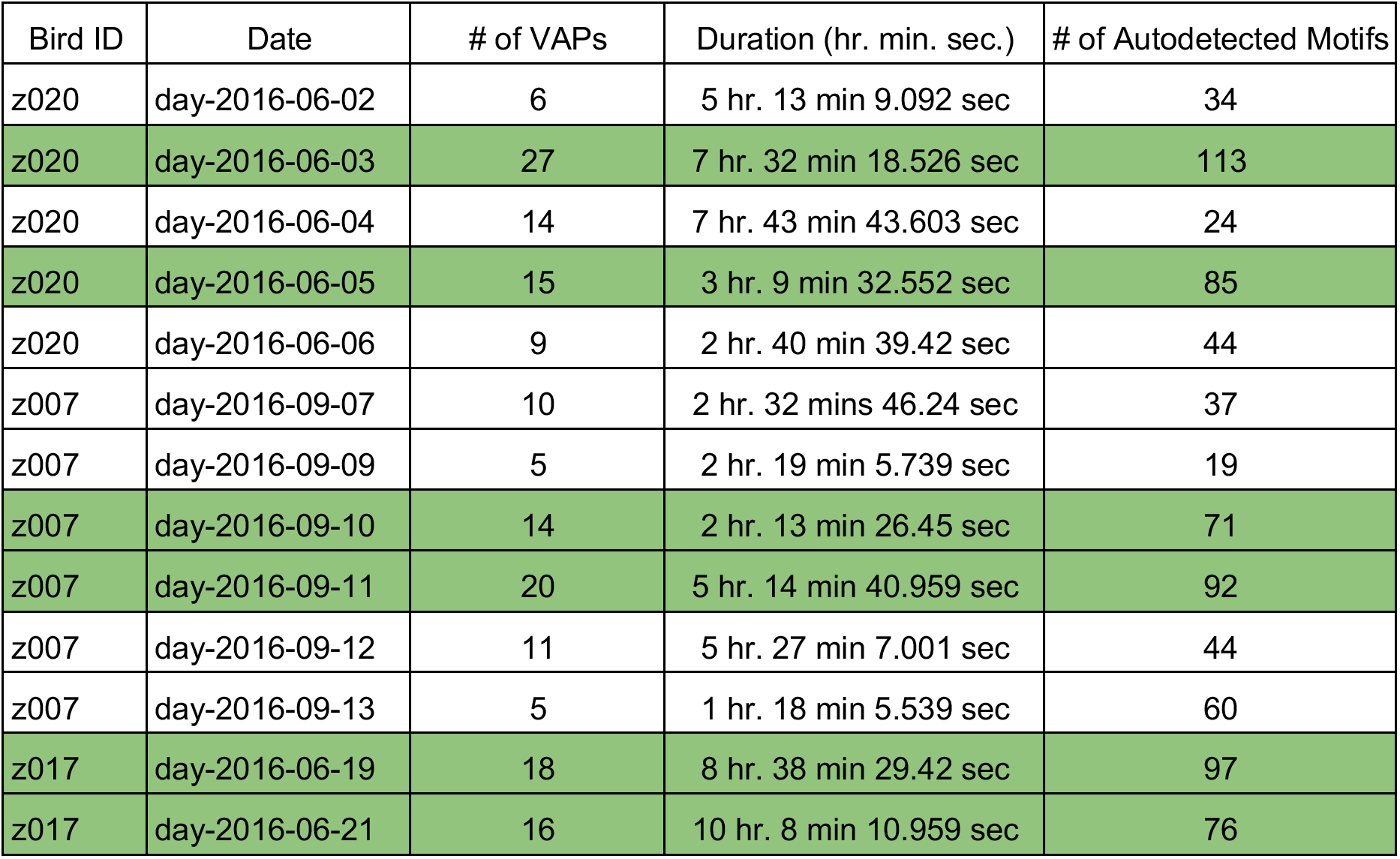
Detailed Description of available labeled data.

| Bird ID | Date | # of VAPs | Duration (hr. min. sec.) | # of Autodetected Motifs |
| --- | --- | --- | --- | --- |
| z020 | day-2016-06-02 | 6 | 5 hr. 13 min 9.092 sec | 34 |
| z020 | day-2016-06-03 | 27 | 7 hr. 32 min 18.526 sec | 113 |
| z020 | day-2016-06-04 | 14 | 7 hr. 43 min 43.603 sec | 24 |
| z020 | day-2016-06-05 | 15 | 3 hr. 9 min 32.552 sec | 85 |
| z020 | day-2016-06-06 | 9 | 2 hr. 40 min 39.42 sec | 44 |
| z007 | day-2016-09-07 | 10 | 2 hr. 32 mins 46.24 sec | 37 |
| z007 | day-2016-09-09 | 5 | 2 hr. 19 min 5.739 sec | 19 |
| z007 | day-2016-09-10 | 14 | 2 hr. 13 min 26.45 sec | 71 |
| z007 | day-2016-09-11 | 20 | 5 hr. 14 min 40.959 sec | 92 |
| z007 | day-2016-09-12 | 11 | 5 hr. 27 min 7.001 sec | 44 |
| z007 | day-2016-09-13 | 5 | 1 hr. 18 min 5.539 sec | 60 |
| z017 | day-2016-06-19 | 18 | 8 hr. 38 min 29.42 sec | 97 |
| z017 | day-2016-06-21 | 16 | 10 hr. 8 min 10.959 sec | 76 |

**S3 Table:**
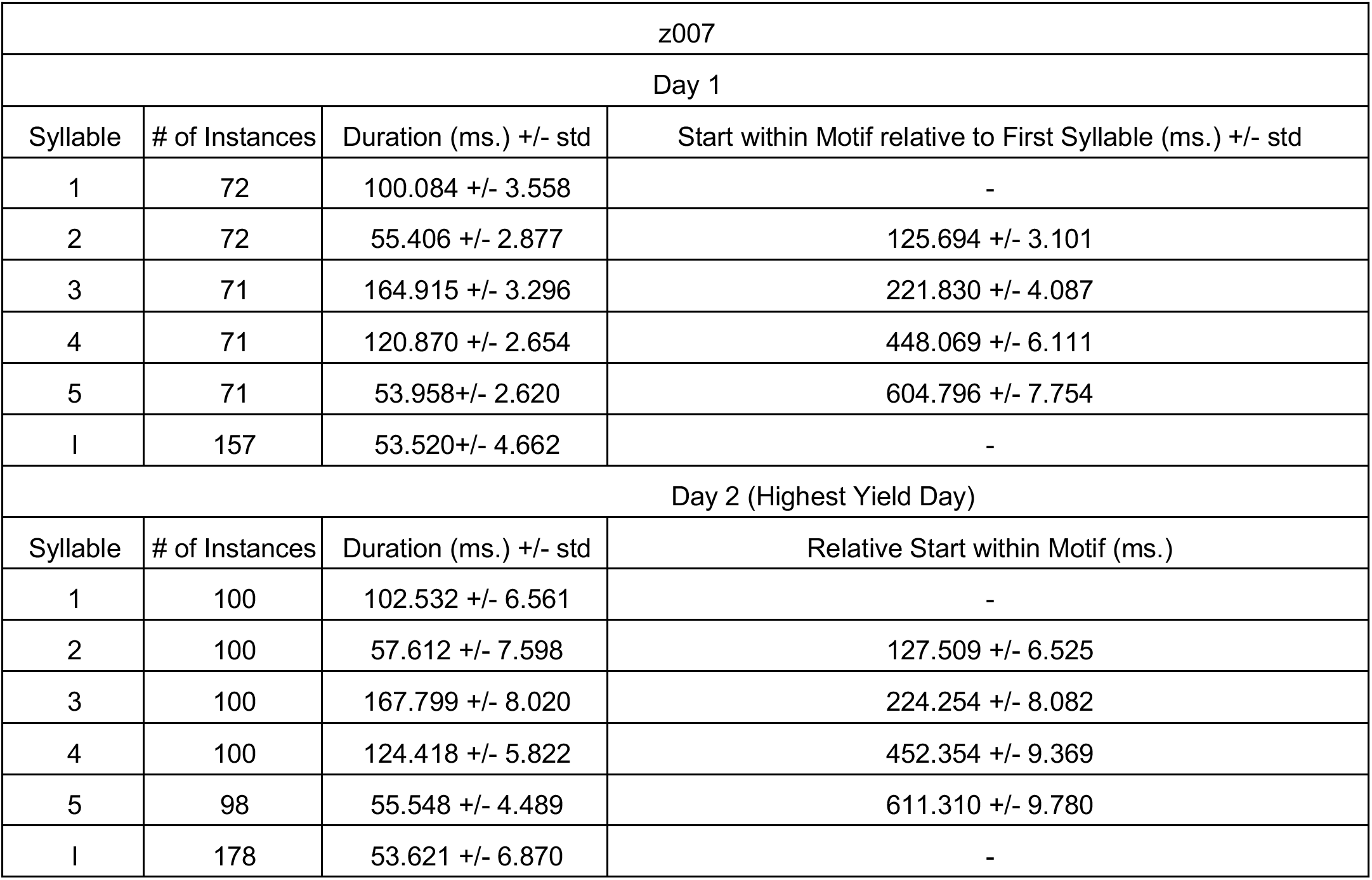
Behavior of Subject z007 during High Yield Days.

| z007 |  |  |  |
| --- | --- | --- | --- |
| Day 1 |  |  |  |
| Syllable | # of Instances | Duration (ms.) +/- std | Start within Motif relative to First Syllable (ms.) +/- std |
| 1 | 72 | 100.084 +/- 3.558 | - |
| 2 | 72 | 55.406 +/- 2.877 | 125.694 +/- 3.101 |
| 3 | 71 | 164.915 +/- 3.296 | 221.830 +/- 4.087 |
| 4 | 71 | 120.870 +/- 2.654 | 448.069 +/- 6.111 |
| 5 | 71 | 53.958 +/- 2.620 | 604.796 +/- 7.754 |
| I | 157 | 53.520 +/- 4.662 | - |
| Day 2 (Highest Yield Day) |  |  |  |
| Syllable | # of Instances | Duration (ms.) +/- std | Relative Start within Motif (ms.) |
| 1 | 100 | 102.532 +/- 6.561 | - |
| 2 | 100 | 57.612 +/- 7.598 | 127.509 +/- 6.525 |
| 3 | 100 | 167.799 +/- 8.020 | 224.254 +/- 8.082 |
| 4 | 100 | 124.418 +/- 5.822 | 452.354 +/- 9.369 |
| 5 | 98 | 55.548 +/- 4.489 | 611.310 +/- 9.780 |
| I | 178 | 53.621 +/- 6.870 | - |

**S4 Table:** Behavior of Subject z020 during High Yield Days.

| z020 |  |  |  |
| --- | --- | --- | --- |
| Day 1 (Highest Yield Day) |  |  |  |
| Syllable | # of Instances | Duration (ms) +/- std | Start within Motif relative to First Syllable (ms) +/- std |
| 1 | 160 | 38.882 +/- 4.716 | - |
| 2 | 160 | 132.902 +/- 8.230 | 65.039 +/- 5.053 |
| 3 | 160 | 49.861 +/- 5.099 | 203.551 +/- 10.469 |
| 4 | 91 | 110.964 +/- 3.476 | 347.219 +/- 23.193 |
| I | 235 | 42.081 +/- 5.792 | - |
| Day 2 |  |  |  |
| Syllable | # of Instances | Duration (ms) +/- std | Relative Start within Motif (ms) +/- std |
| 1 | 109 | 39.887 +/- 6.139 | - |
| 2 | 109 | 132.925 +/- 4.865 | 67.670 +/- 5.01 |
| 3 | 109 | 50.257 +/- 5.011 | 206.310 +/- 4.985 |
| 4 | 75 | 111.741 +/- 6.112 | 348.720 +/- 12.329 |
| I | 191 | 46.922 +/- 9.378 | - |

**S5 Table:** Behavior of Subject z017 during High Yield Days.

| z017 |  |  |  |
| --- | --- | --- | --- |
| Day 1 (Highest Yield Day) |  |  |  |
| Syllable | # of Instances | Duration (ms) +/- std | Start within Motif relative to First Syllable (ms) +/- std |
| 1 | 97 | 50.692 +/- 5.341 | - |
| 2 | 97 | 57.389 +/- 6.629 | 73.109 +/- 5.061 |
| 3 | 97 | 117.752 +/- 7.757 | 157.192 +/- 7.368 |
| 4 | 97 | 106.304 +/- 5.832 | 307.916 +/- 9.093 |
| 5 | 97 | 127.465 +/- 4.870 | 420.749 +/- 9.343 |
| 6 | 97 | 154.392 +/- 6.617 | 591.343 +/- 13.956 |
| 7 | 52 | 72.894 +/- 8.067 | 809.046 +/- 21.541 |
| Day 2 |  |  |  |
| Syllable | # of Instances | Duration (ms) +/- std | Relative Start within Motif (ms) +/- std |
| 1 | 84 | 49.162 +/- 6.525 | - |
| 2 | 84 | 51.217 +/- 9.207 | 75.058 +/- 6.447 |
| 3 | 86 | 114.807 +/- 15.198 | 155.762 +/- 12.082 |
| 4 | 83 | 106.109 +/- 8.912 | 306.191 +/- 18.28 |
| 5 | 82 | 127.825 +/- 11.710 | 420.302 +/- 22.832 |
| 6 | 66 | 154.634 +/- 11.803 | 584.710 +/- 14.729 |
| 7 | 41 | 68.696 +/- 8.431 | 803.838 +/- 26.629 |

**S6 Table:** Characteristic Increases in Power in Narrowband Frequencies during Song Across Channels. The percentage of channels who’s distribution of power values were greater during periods of vocal activity than during silence. To determine statistical significance between the two distributions the one sided z-test was used with each frequency and channel pair. The percentages shown are the number of good channels that still were significant after using the using the Benjamini-Hochberg False Discovery Rate (p<0.05 and q<0.05).

| Bird_ID | Date | Percentage of Channels with Increased Power During Vocally Active Periods<br>(After FDR Correction) |  |  |  |  |  |
| --- | --- | --- | --- | --- | --- | --- | --- |
|  |  | 4-8 Hz | 8-12 Hz | 25-35 Hz | 35-50 Hz | 50-70 Hz | 80-200 Hz |
| z020 | day-2016-06-03 | 100.00% | 100.00% | 100.00% | 100.00% | 100.00% | 100.00% |
| z020 | day-2016-06-05 | 53.33% | 26.66% | 26.66% | 26.66% | 40.00% | 26.66% |
| z007 | day-2016-09-10 | 70.00% | 76.66% | 86.66% | 100.00% | 100.00% | 100.00% |
| z007 | day-2016-09-11 | 73.33% | 80.00% | 100.00% | 100.00% | 100.00% | 100.00% |
| z017 | day-2016-06-19 | 0.00% | 6.25% | 62.50% | 68.75% | 87.50% | 100.00% |
| z017 | day-2016-06-21 | 0.00% | 6.25% | 81.25% | 81.25% | 100.00% | 100.00% |

**S7 Table:** Characteristic Decreases in Power in Narrowband Frequencies during Song Across Channels. The percentage of channels who’s distribution of power values were smaller during periods of vocal activity than during silence. To determine statistical significance between the two distributions the one sided z-test was used with each frequency and channel pair. The percentages shown are the number of good channels that still were significant after using the using the Benjamini-Hochberg False Discovery Rate (p<0.05 and q<0.05).

| Bird_ID | Date | Percentage of Channels with Decreased Power During Vocally Active Periods<br>(After FDR Correction) |  |  |  |  |  |
| --- | --- | --- | --- | --- | --- | --- | --- |
|  |  | 4-8 Hz | 8-12 Hz | 25-35 Hz | 35-50 Hz | 50-70 Hz | 80-200 Hz |
| z020 | day-2016-06-03 | 0.00% | 0.00% | 0.00% | 0.00% | 0.00% | 0.00% |
| z020 | day-2016-06-05 | 40.00% | 53.33% | 40.00% | 60.00% | 46.60% | 60.00% |
| z007 | day-2016-09-10 | 30.00% | 19.99% | 3.33% | 0.00% | 0.00% | 0.00% |
| z007 | day-2016-09-11 | 26.66% | 19.99% | 0.00% | 0.00% | 0.00% | 0.00% |
| z017 | day-2016-06-19 | 100.00% | 93.75% | 31.25% | 31.25% | 0.00% | 0.00% |
| z017 | day-2016-06-21 | 100.00% | 87.75% | 6.25% | 0.00% | 0.00% | 0.00% |

**S8 Table:** Bin Width Hyperparameter Search Results.

| Bird ID | Date | Best Bin Width (ms) |  |  |  |  |
| --- | --- | --- | --- | --- | --- | --- |
|  |  | 4-8 Hz | 8-12 Hz | 25-35 Hz | 35-50 Hz | 50-70 Hz |
| z020 | day-2016-06-03 | 90 | 185 | 90 | 85 | 90 |
| z020 | day-2016-06-05 | 190 | 125 | 35 | 160 | 70 |
| z007 | day-2016-09-10 | 170 | 75 | 90 | 25 | 30 |
| z007 | day-2016-09-11 | 195 | 170 | 70 | 20 | 70 |
| z017 | day-2016-06-19 | 175 | 170 | 85 | 50 | 35 |
| z017 | day-2016-06-21 | 150 | 170 | 70 | 60 | 40 |

**S9 Table:** Offset Hyperparameter Search Results.

| Bird ID | Date | Best Offset (ms) |  |  |  |  |
| --- | --- | --- | --- | --- | --- | --- |
|  |  | 4-8 Hz | 8-12 Hz | 25-35 Hz | 35-50 Hz | 50-70 Hz |
| z020 | day-2016-06-03 | -10 | -5 | 0 | 0 | 0 |
| z020 | day-2016-06-05 | -25 | -5 | 0 | -20 | 0 |
| z007 | day-2016-09-10 | -35 | -5 | -10 | -10 | 0 |
| z007 | day-2016-09-11 | -5 | -5 | -15 | 0 | -15 |
| z017 | day-2016-06-19 | -50 | 0 | -10 | -5 | -5 |
| z017 | day-2016-06-21 | -10 | -15 | -30 | 0 | 0 |

**S1 Fig:**
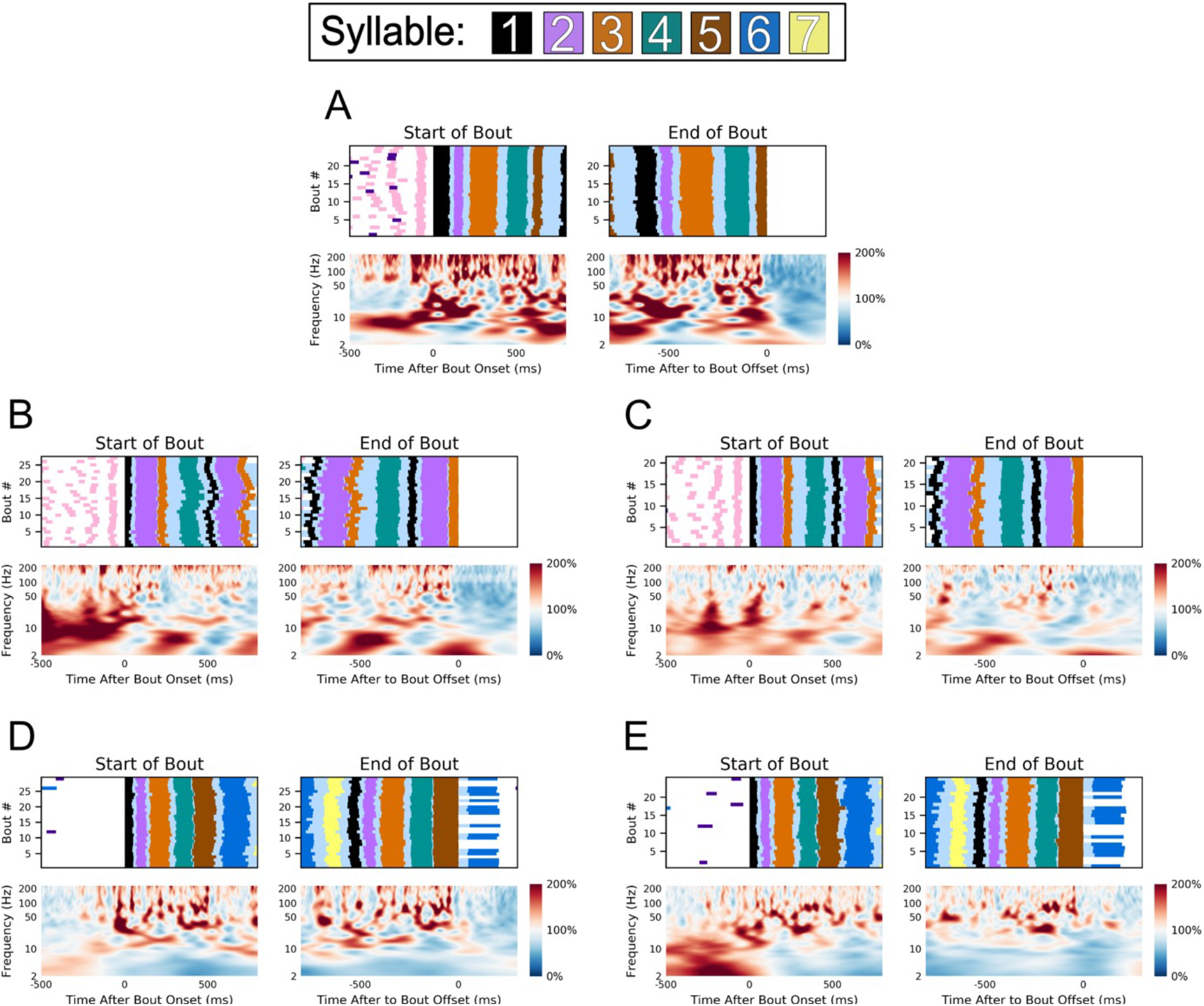
Song correlated changes in LFP power across subjects and days. Averaged spectrotemporal power activity (see **Methods**) aligned to the start of the first motif in the bout, left, and the last motif in the bout, right, for each recording day not shown in Fig 3. Above all results is a behavioral raster showing the time course of the behavior being averaged. (A) The averaged results for the second highest yielding day, designated Day 1, for z007 (n=25 Bouts). The other subjects results are show as follows; (B) z020’s first high yield day (n=29 Bouts), (C) z020’s second high yield day (n=25 Bouts), (D) z017’s first high yield day (n=27 Bouts), and (E) z017’s second high yield day (n=21 Bouts). Because z017 would end its bout on either syllable ‘5’ or ‘6’, the end of the bout was aligned to syllable ‘5’. No dynamic-time warping was used. To ensure the start and end of the bout are unique time periods, only bouts with more than one motif in duration were used. Behaviorally inconsistent bouts were excluded for clarity of visualization, however results are consistent when including them in the analysis.

**S2 Fig:**
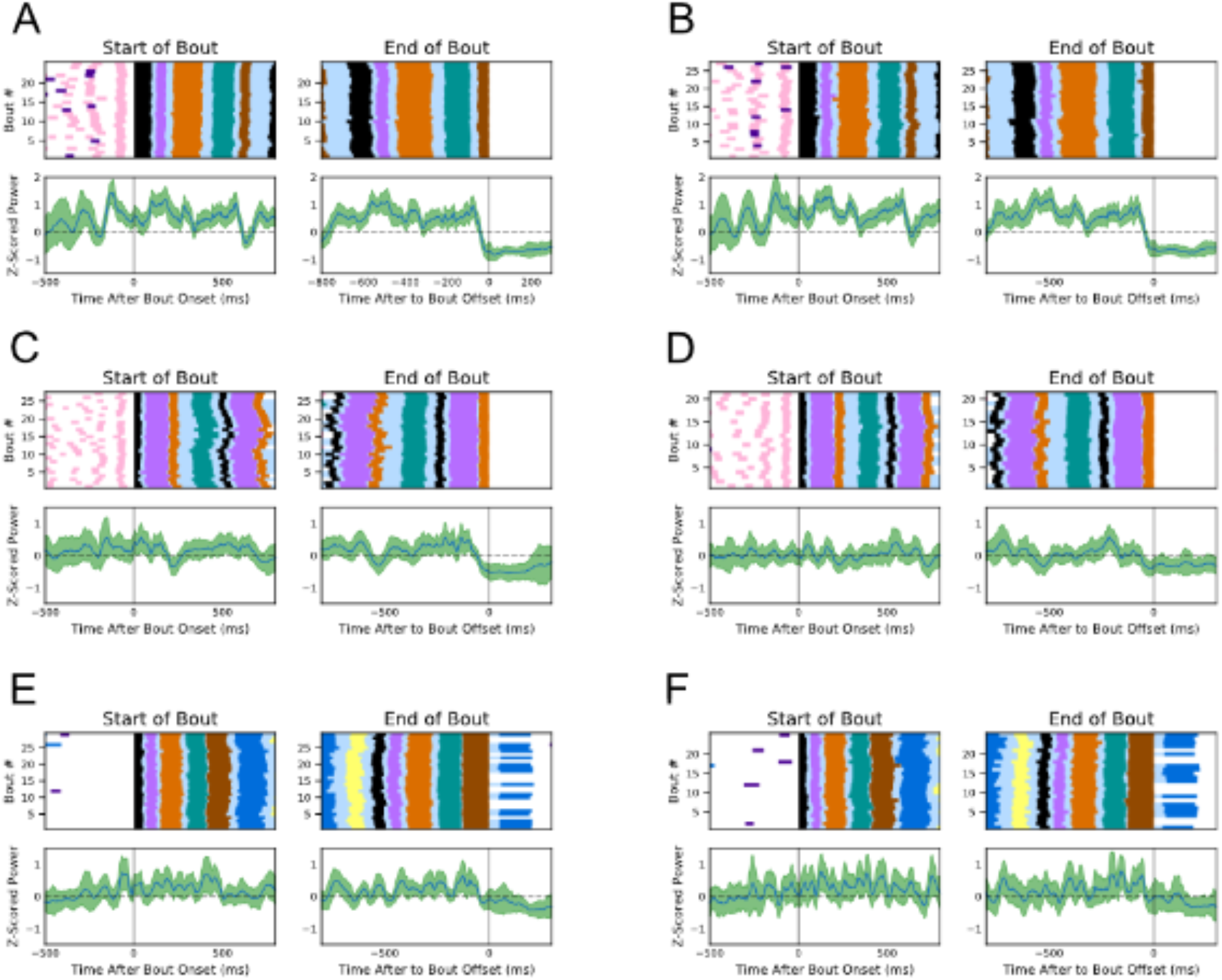
Song correlated changes in power of 50-200 Hz band across days and subjects. Z-Scored Changes in power of the 50-200 Hz Band aligned to the start of the first motif in the bout, left, and the last motif in the bout, right, for each high yield recording day. Black Traces in each subpanel is the mean and the green shading is the standard deviation. Above all results is a behavioral raster showing the time course of the behavior being averaged. (A) The results of the first high yield Day of z007 (n=27 Bouts) (B) The results for the second high yield day for z007 (n=25 Bouts). The other subjects results are show as follows; (C) z020’s first high yield day (n=29 Bouts), (D) z020’s second high yield day (n=25 Bouts), (E) z017’s first high yield day (n=27 Bouts), and (F) z017’s second high yield day (n=21 Bouts). Because z017 would end its bout on either syllable ‘5’ or ‘6’, the end of the bout was aligned to syllable ‘5’. No dynamic-time warping was used. To ensure the start and end of the bout are unique time periods, only bouts with more than one motif in duration were used. Behaviorally inconsistent bouts were excluded for clarity of visualization, however results are consistent when including them in the analysis.

**S3 Fig:**
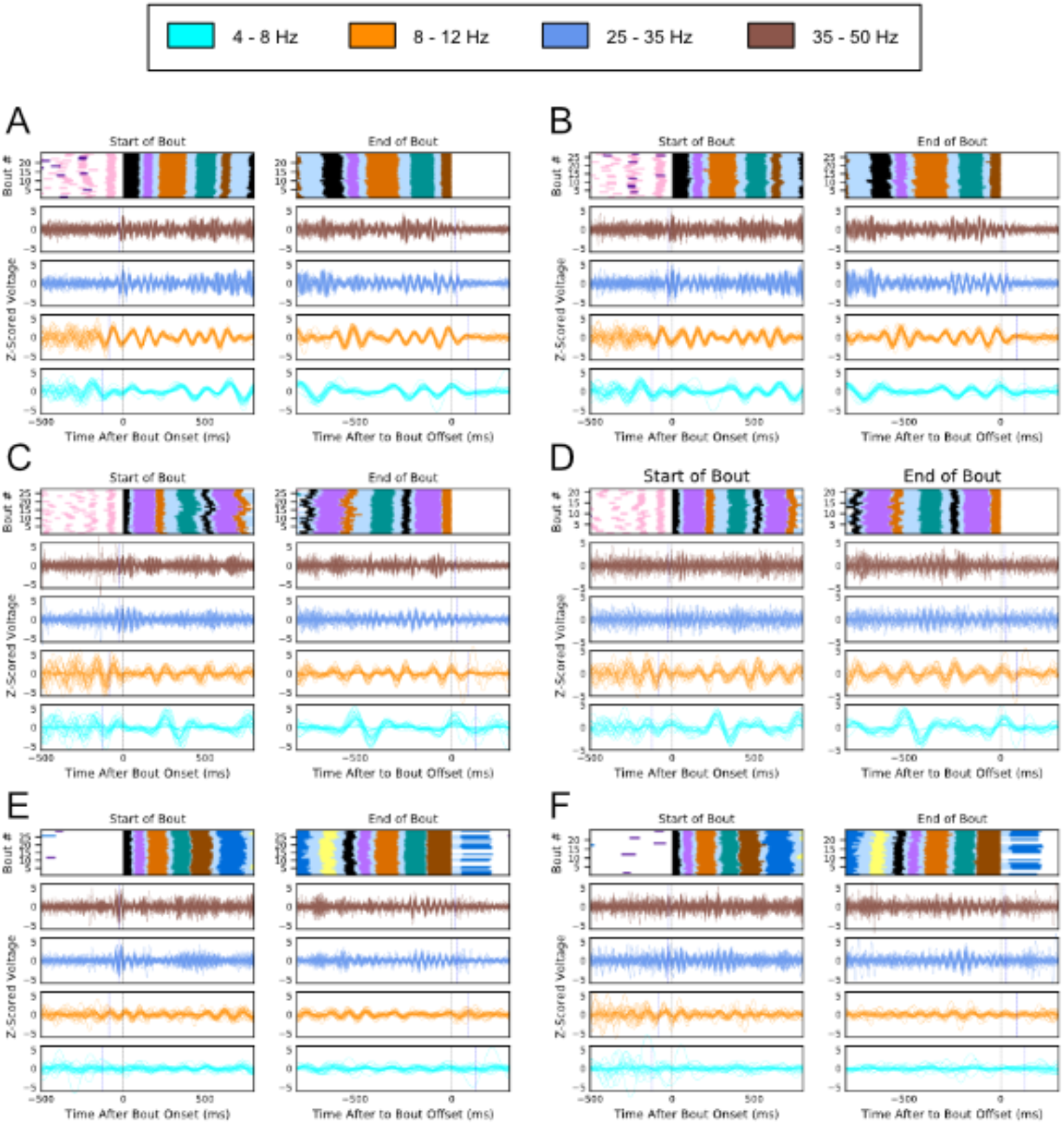
Stereotyped song correlated rhythmic changes in LFP across days and subjects (single trial). Single Trial z-scored LFP traces of four Narrowband frequency bands aligned to the start of the first motif in the bout, left, and the last motif in the bout, right, for each high yield recording day. Each Trace is colored its respective narrowband frequency. Above all results is a behavioral raster showing the time course of the behaviors being show. The black line in each row below the behavior shows the time point the trials are aligned to, and the blue line denotes the time prior to (left) or after (right) one full cycle of the highest frequency in the narrowband frequency. (A) The results of the first high yield Day of z007 (n=27 Bouts) (B) The results for the second high yield day for z007 (n=25 Bouts). The other subjects results are show as follows; (C) z020’s first high yield day (n=29 Bouts), (D) z020’s second high yield day (n=25 Bouts), (E) z017’s first high yield day (n=27 Bouts), and (F) z017’s second high yield day (n=21 Bouts). Because z017 would end its bout on either syllable ‘5’ or ‘6’, the end of the bout was aligned to syllable ‘5’. No dynamic-time warping was used. To ensure the start and end of the bout are unique time periods, only bouts with more than one motif in duration were used. Behaviorally inconsistent bouts were excluded for clarity of visualization, however results are consistent when including them in the analysis.

**S4 Fig:**
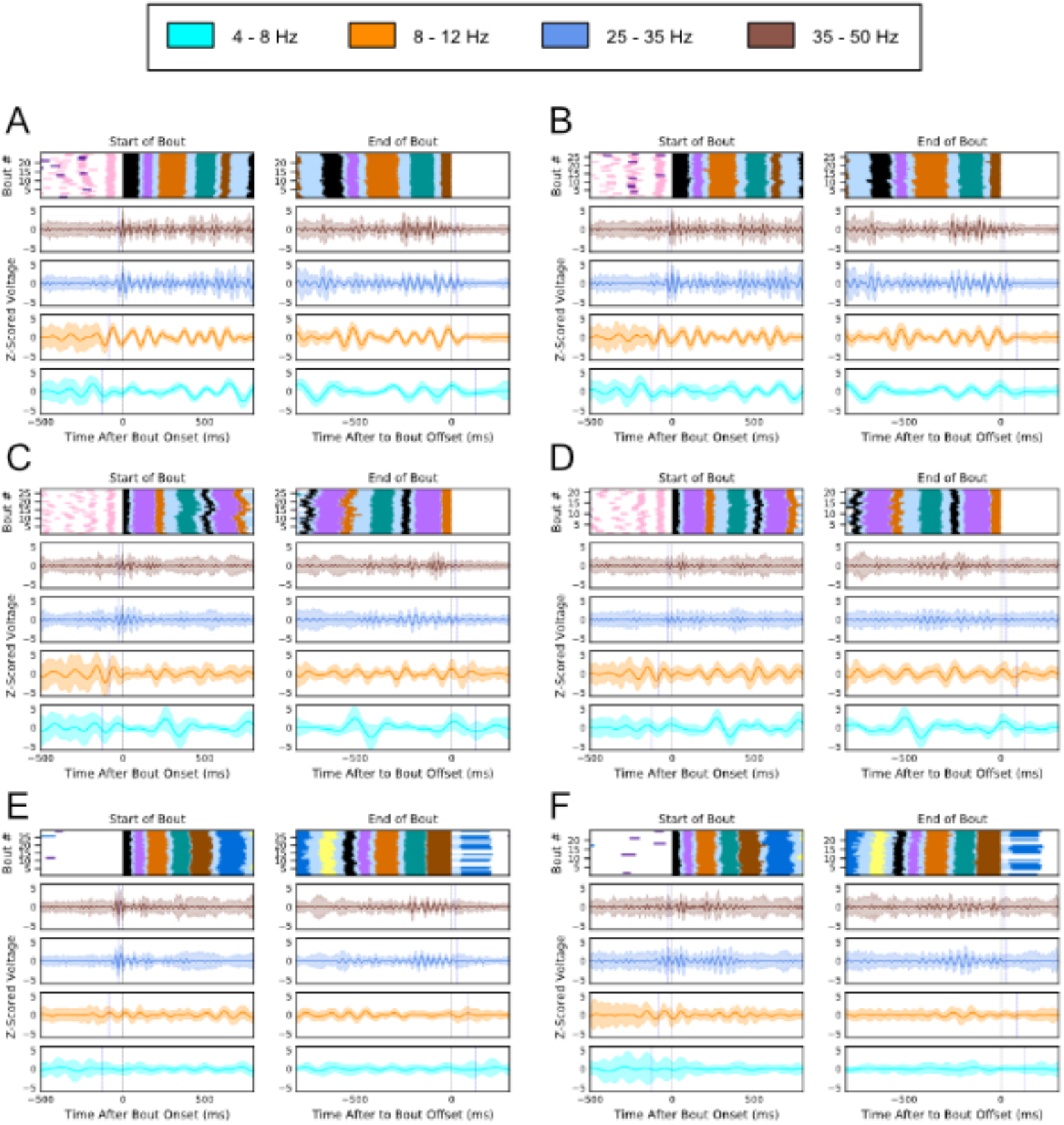
Stereotyped song correlated rhythmic changes in LFP across days and subjects (mean and standard deviation). The Mean and standard deviation of the z-scored LFP traces of four Narrowband frequency bands aligned to the start of the first motif in the bout, left, and the last motif in the bout, right, for each high yield recording day. Each row is colored its respective narrowband frequency. Above all results is a behavioral raster showing the time course of the behaviors being show. The black line in each row below the behavior shows the time point the trials are aligned to, and the blue line denotes the time prior to (left) or after (right) one full cycle of the highest frequency in the narrowband frequency. (A) The results of the first high yield Day of z007 (n=27 Bouts) (B) The results for the second high yield day for z007 (n=25 Bouts). The other subjects results are show as follows; (C) z020’s first high yield day (n=29 Bouts), (D) z020’s second high yield day (n=25 Bouts), (E) z017’s first high yield day (n=27 Bouts), and (F) z017’s second high yield day (n=21 Bouts). Because z017 would end its bout on either syllable ‘5’ or ‘6’, the end of the bout was aligned to syllable ‘5’. No dynamic-time warping was used. To ensure the start and end of the bout are unique time periods, only bouts with more than one motif in duration were used. Behaviorally inconsistent bouts were excluded for clarity of visualization, however results are consistent when including them in the analysis.

**S5 Fig:**
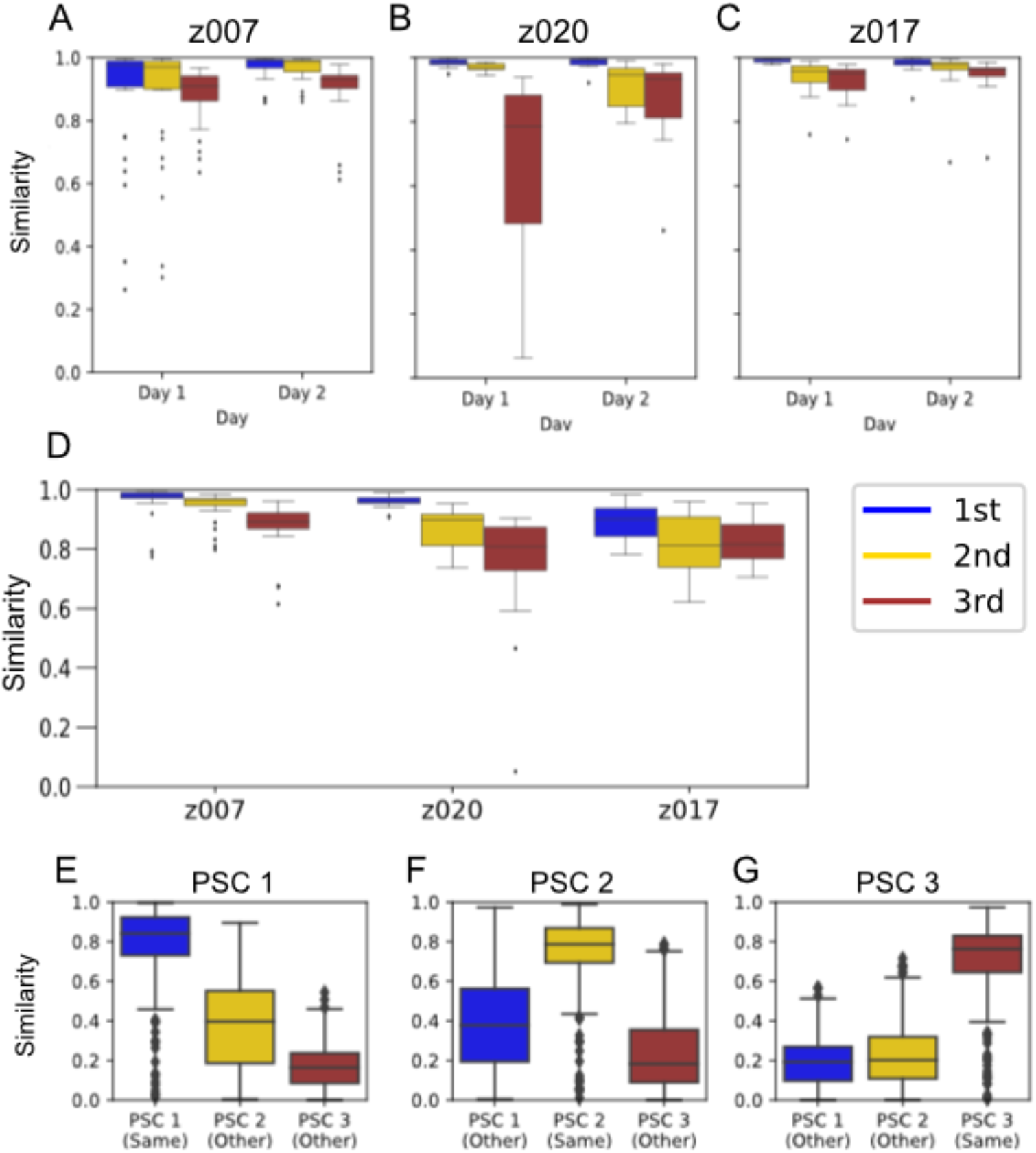
Similarity of principal spectral components across channels, days and subjects. Boxplot of the distribution of Cosine Similarity Metric values between a template, which is created by taking the mean across the sign aligned PSC of all channels for a specific PSC. The cosine similarity Matrix ranges from 1 and −1, however the absolute value of the metric is shown (see Methods). (A) The cosine similarity of each channel’s PSC to the same recording day’s template PSC for both high yield Days subject z007. (B) The cosine similarity of each channel’s PSC to the same recording day’s template PSC for both high yield Days subject z020. (C) The cosine similarity of each channel’s PSC to the same recording day’s template PSC for both high yield Days subject z017. (D) The cosine similarity of each channel’s PSC from the second high yield day with the template of the PSC from the first high yield day. (E) All templates for PSC 1 compared either with the PSC 1 for the other two birds, same, or the PSC for one of the other PSCs for the other two birds, other. (F) All templates for PSC 2 compared either with the PSC 2 for the other two birds, same, or the PSC for one of the other PSCs for the other two birds, other. (E) All templates for PSC 3 compared either with the PS C3 for the other two birds, same, or the PSC for one of the other PSCs for the other two birds, other.

**S6 Fig:**
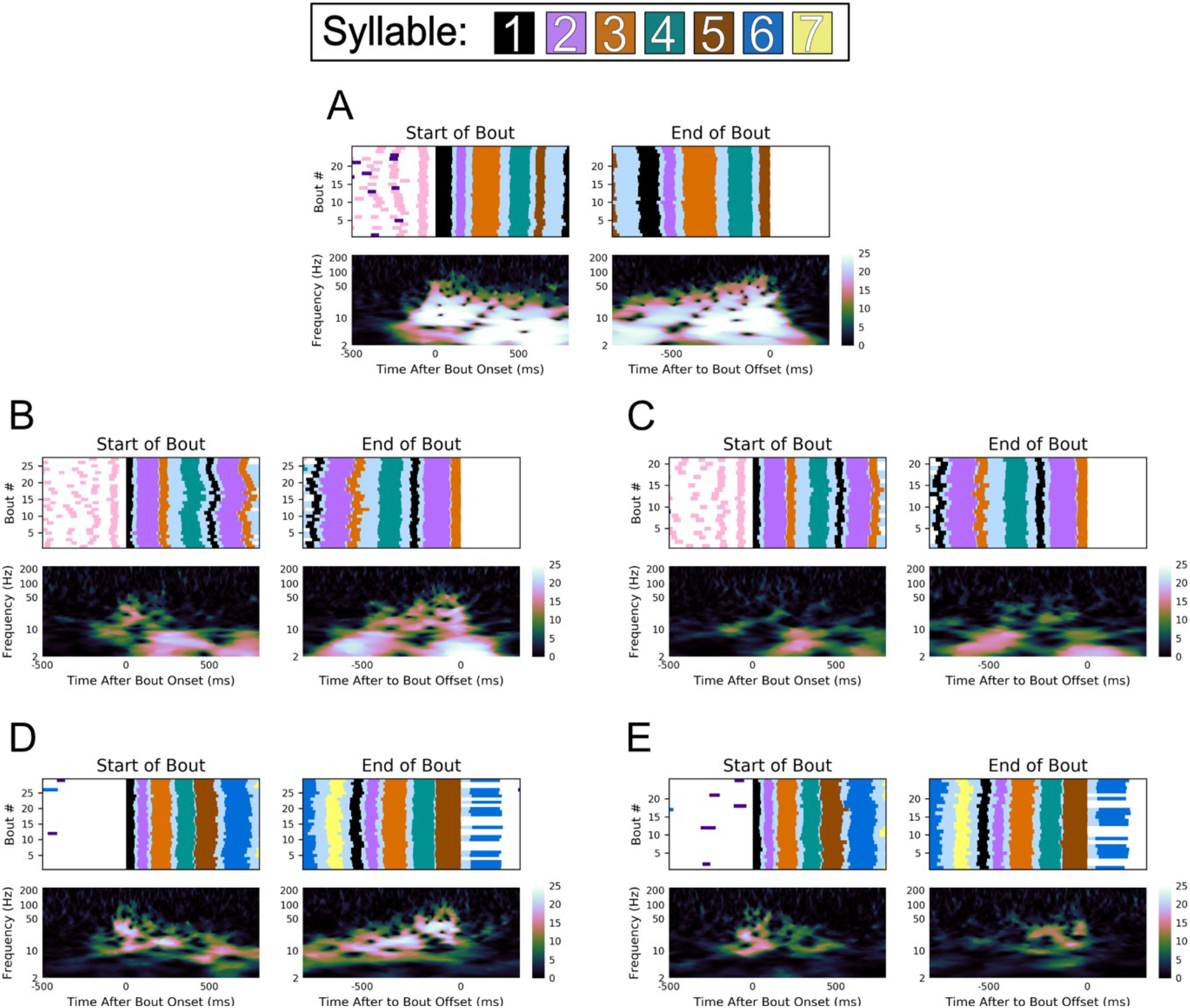
Inter-trial phase coherence of LFP phase during production of learned sequences across subjects and days. ITPC of LFP aligned to the start of the first motif in the bout, left, and the last motif in the bout, right, for each recording day not shown in Fig 3. Above all results is a Behavioral raster showing the time course of the behavior being averaged. (A) The averaged results for the second highest yielding day, designated Day 1, for z007 (n=25 bouts). The other subjects results are show as follows; (B) z020’s first high yield day (n=29 bouts), (C) z020’s second high yield day (n=25 bouts), (D) z017’s first high yield day (n=27 bouts), and (E) z017’s second high yield day (n=21 bouts). Because z017 would end its bout on either syllable ‘5’ or ‘6’, the end of the bout was aligned to syllable ‘5’. No dynamic-time warping was used. To ensure the start and end of the bout are unique time periods, only bouts with more than one motif in duration were used. Behaviorally inconsistent bouts were excluded for clarity of visualization, however results are consistent when including them in calculating the ITPC. (p<0.006 for all Z > 5 for all subjects and days).

**S7 Fig:**
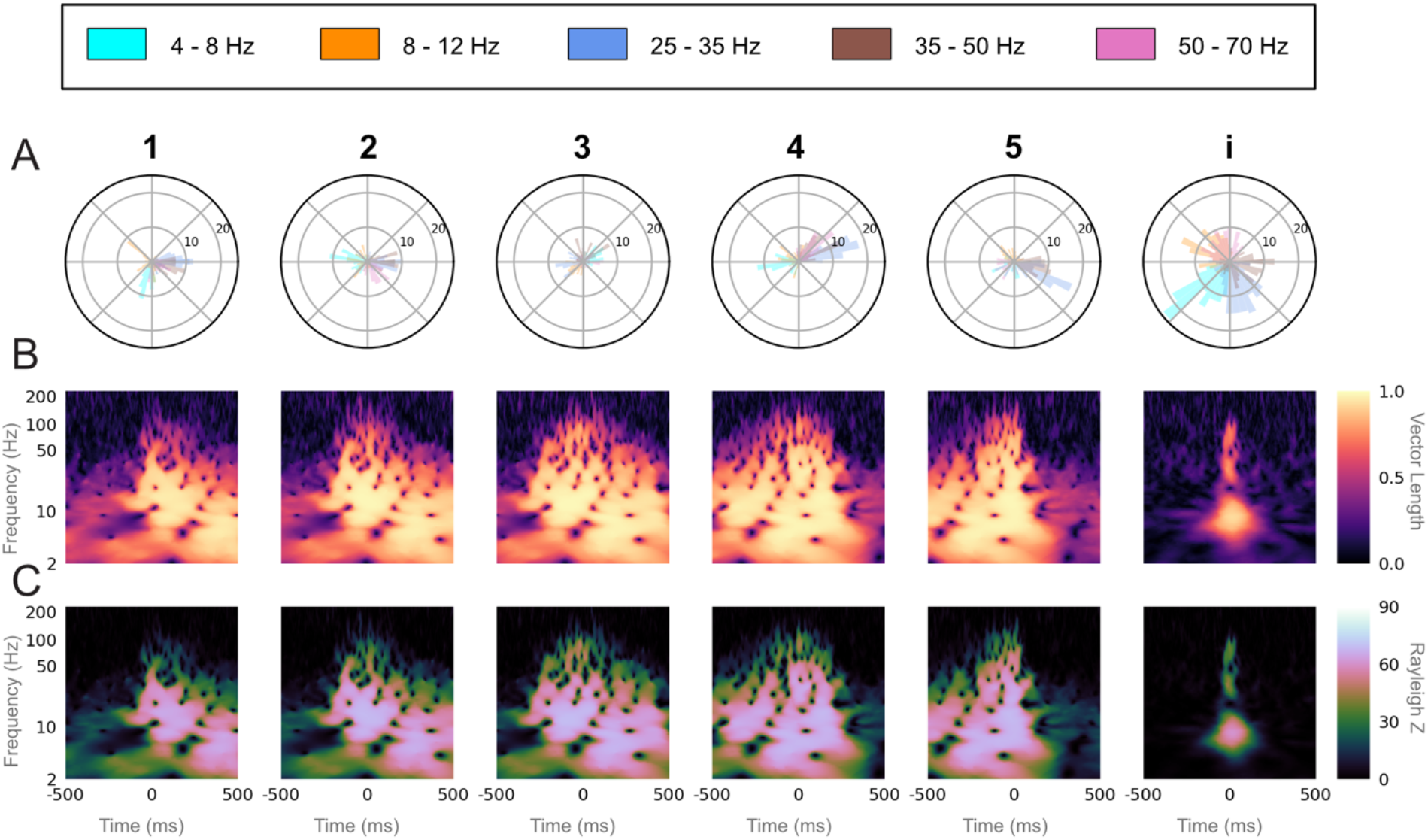
Unique LFP phase preferences at each syllable onset (z007, day 1). (A) Polar histogram of the phase for each LFP frequency band at the labeled start of all instances of a given syllable or the introductory note over the course of one day (Day 1), for one bird (z007). The number of instances (n=71) are equal for all syllables and the introductory note, and is set by the syllable class with the fewest renditions. (B) ITPC resultant vector length for each frequency over time relative to the labeled start of each syllable or introductory note (0 ms) over the same instances as in (A). (C) Rayleigh Z-statistic of the ITPC over the same time and frequencies as (B). (p<0.007 for all Z > 5 for all syllables and the introductory note).

**S8 Fig:**
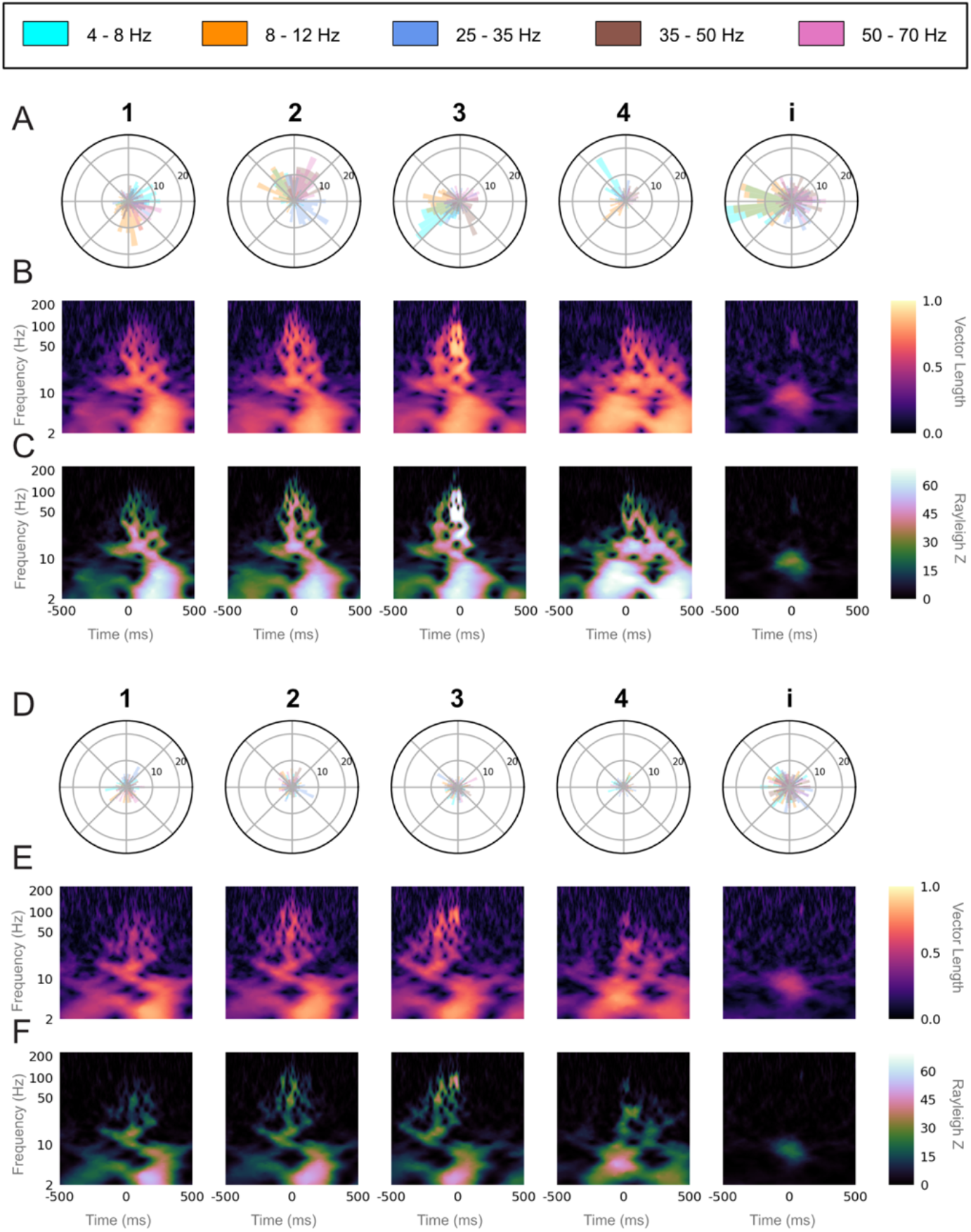
Unique LFP phase preferences at each syllable onset (z020). (A) Polar histogram of the phase for each LFP frequency band at the labeled start of all instances of a given syllable or the introductory note over the course of one day (Day 1), for one bird (z020). The number of instances (n=91) are equal for all syllables and the introductory note, and is set by the syllable class with the fewest renditions. (B) ITPC resultant vector length for each frequency over time relative to the labeled start of each syllable or introductory note (0 ms) over the same instances as in (A). (C) Rayleigh Z-statistic of the ITPC over the same time and frequencies as (B). (D) Polar histogram of the phase for each LFP frequency band at the labeled start of all instances of a given syllable over the course of one day (Day 2), for one bird (z020). The number of instances (n=75) are equal for all syllables and the introductory note, and is set by the syllable class with the fewest renditions. (E) ITPC resultant vector length for each frequency over time relative to the labeled start of each syllable (0 ms) over the same instances as in (D). (F) Rayleigh Z-statistic of the ITPC over the same time and frequencies as (E). (p<0.007 for all Z > 5 for all syllables and the introductory note for both days).

**S9 Fig:**
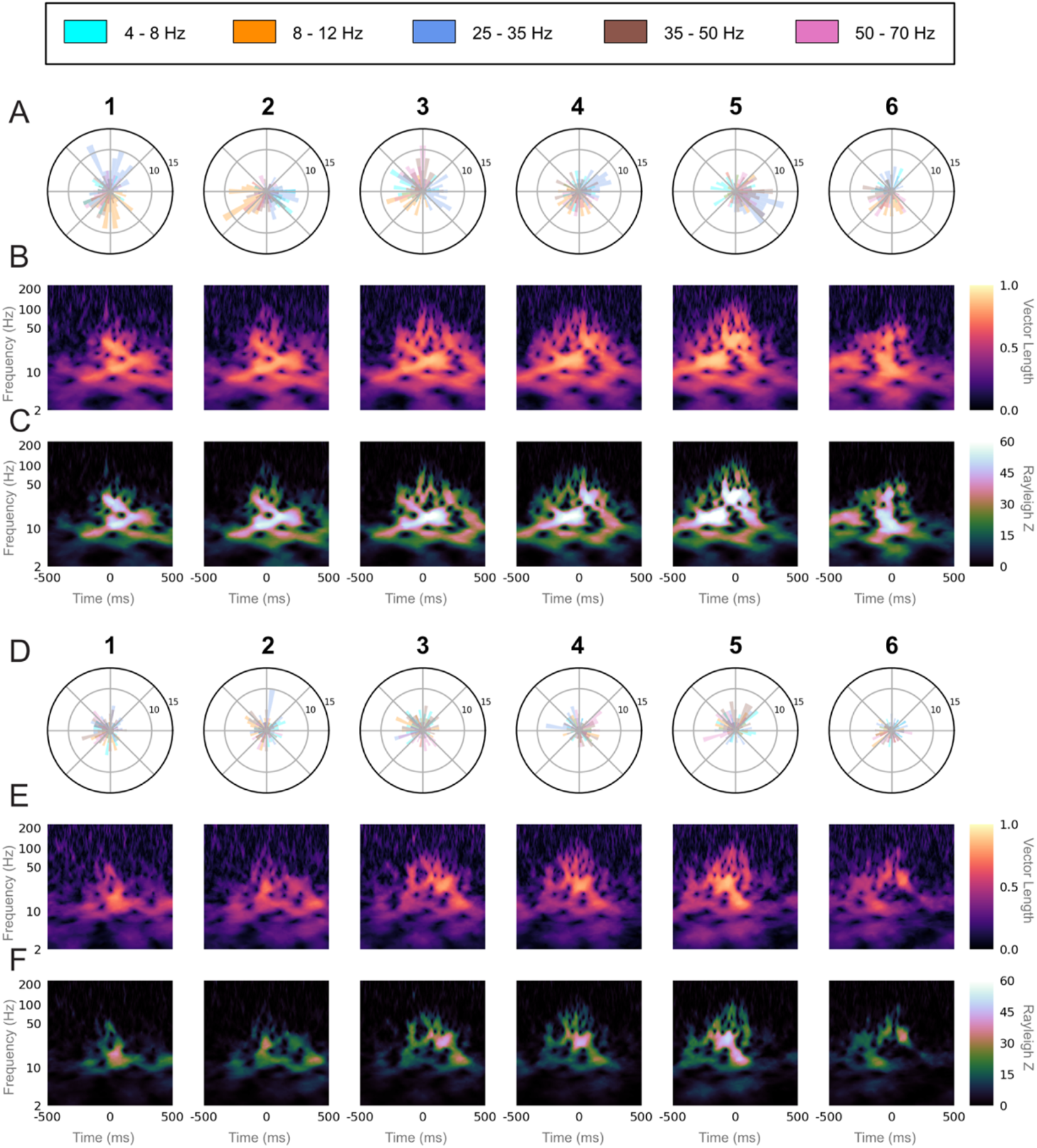
Unique LFP phase preferences at each syllable onset (z007). (A) Polar histogram of the phase for each LFP frequency band at the labeled start of all instances of a given syllable over the course of one day (Day 1), for one bird (z020). The number of instances (n=97) are equal for all syllables, and set by the syllable class with the fewest renditions. (B) ITPC resultant vector length for each frequency over time relative to the labeled start of each syllable (0 ms) over the same instances as in (A). (C) Rayleigh Z-statistic of the ITPC over the same time and frequencies as (B). (D) Polar histogram of the phase for each LFP frequency band at the labeled start of all instances of a given syllable over the course of one day (Day 2), for one bird (z020). The number of instances (n=82) are equal for all syllables, and set by the syllable class with the fewest renditions. (E) ITPC resultant vector length for each frequency over time relative to the labeled start of each syllable (0 ms) over the same instances as in (D). (F) Rayleigh Z-statistic of the ITPC over the same time and frequencies as (E). (p<0.007 for all Z > 5 for all syllables for both days).

**S10 Fig:**
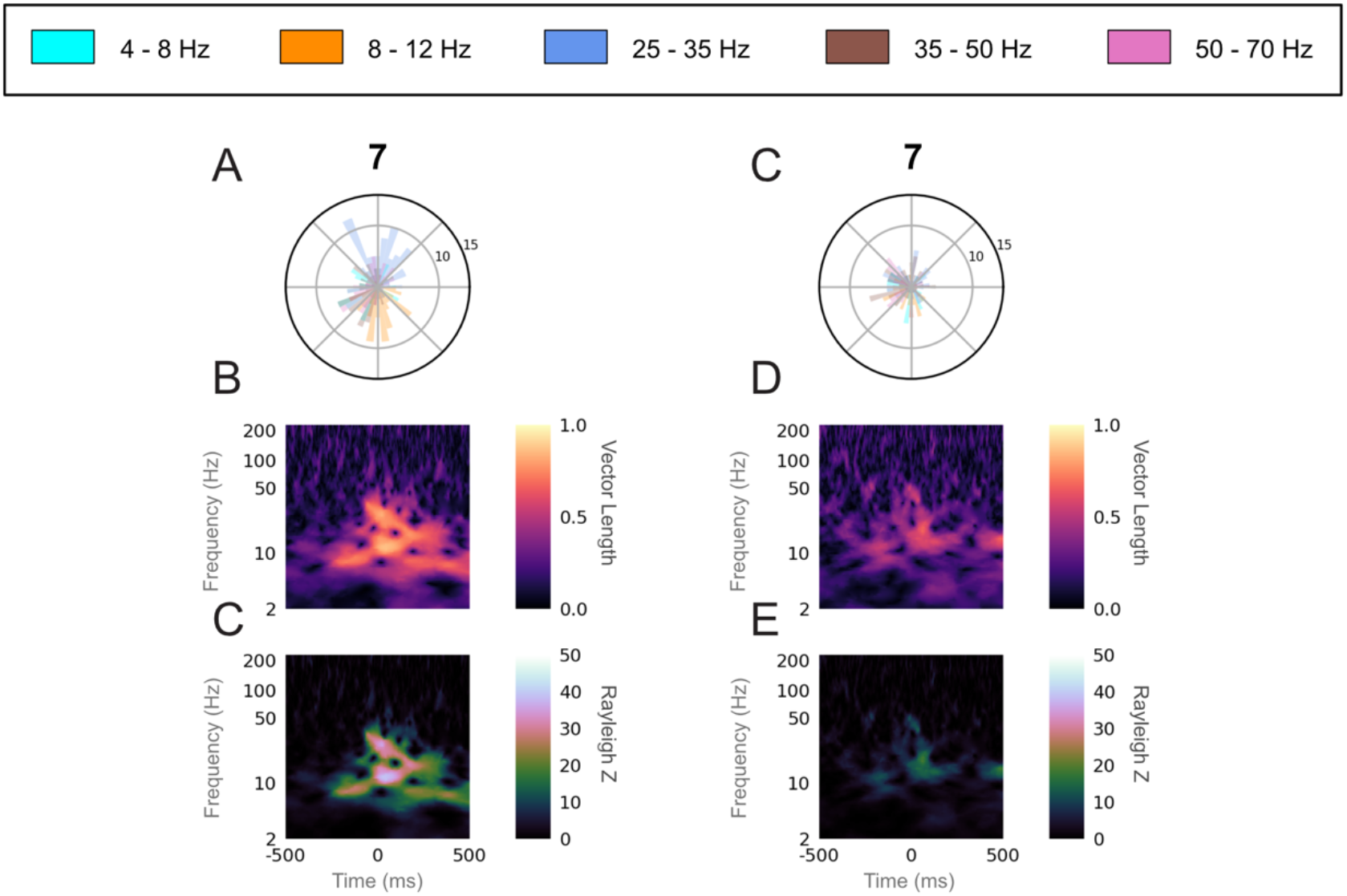
Unique LFP phase preferences for sparsely used intra-motif note onset (z017). (A) Polar histogram of the phase for each LFP frequency band at the labeled start of all instances (n=52) of syllable 7 over the course of one day (Day 1), for bird z017. (B) ITPC resultant vector length for each frequency over time relative to the labeled start of syllable 7 (0 ms) over the same instances as in (A). (C) Rayleigh Z-statistic of the ITPC over the same time and frequencies as (B). (D) Polar histogram of the phase for each LFP frequency band at the labeled start of all instances (n=41) of syllable 7 over the course of one day (Day 2), for bird z017. (E) ITPC resultant vector length for each frequency over time relative to the labeled start of syllable 7 (0 ms) over the same instances as in (D). (F) Rayleigh Z-statistic of the ITPC over the same time and frequencies as (E). (p<0.007 for all Z > 5 for both syllables).

**S11 Fig:**
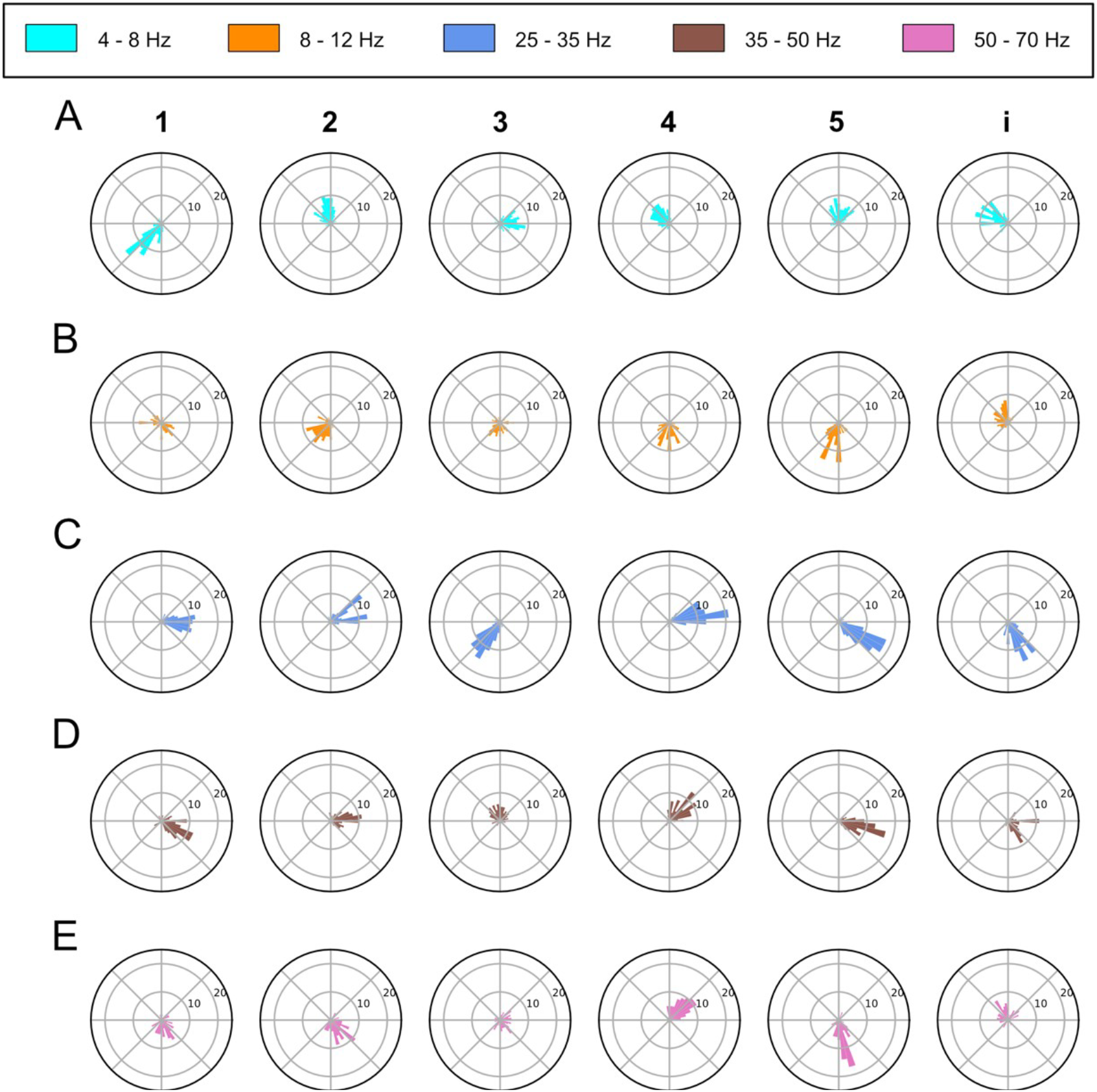
Detailed view of the syllable phase preference to syllable onset for z007. Detailed rendering of the phase preference to syllable onset for z007 shown in Fig 6A. Each Row is a different vocalization type which includes the five syllables of the motif and the introductory note. Each column is a different frequency band and is organized top to bottom from least to greatest. As such they are the polar plots for the (A) 4-8 Hz band, (B) 8-12 Hz band, (C) 25-35 Hz band, (D) 35-50 Hz band, and (E) the 50-70 Hz band. The number of instances have been balanced to match the class with the least number of instances (n=100) for each class.

**S12 Fig:**
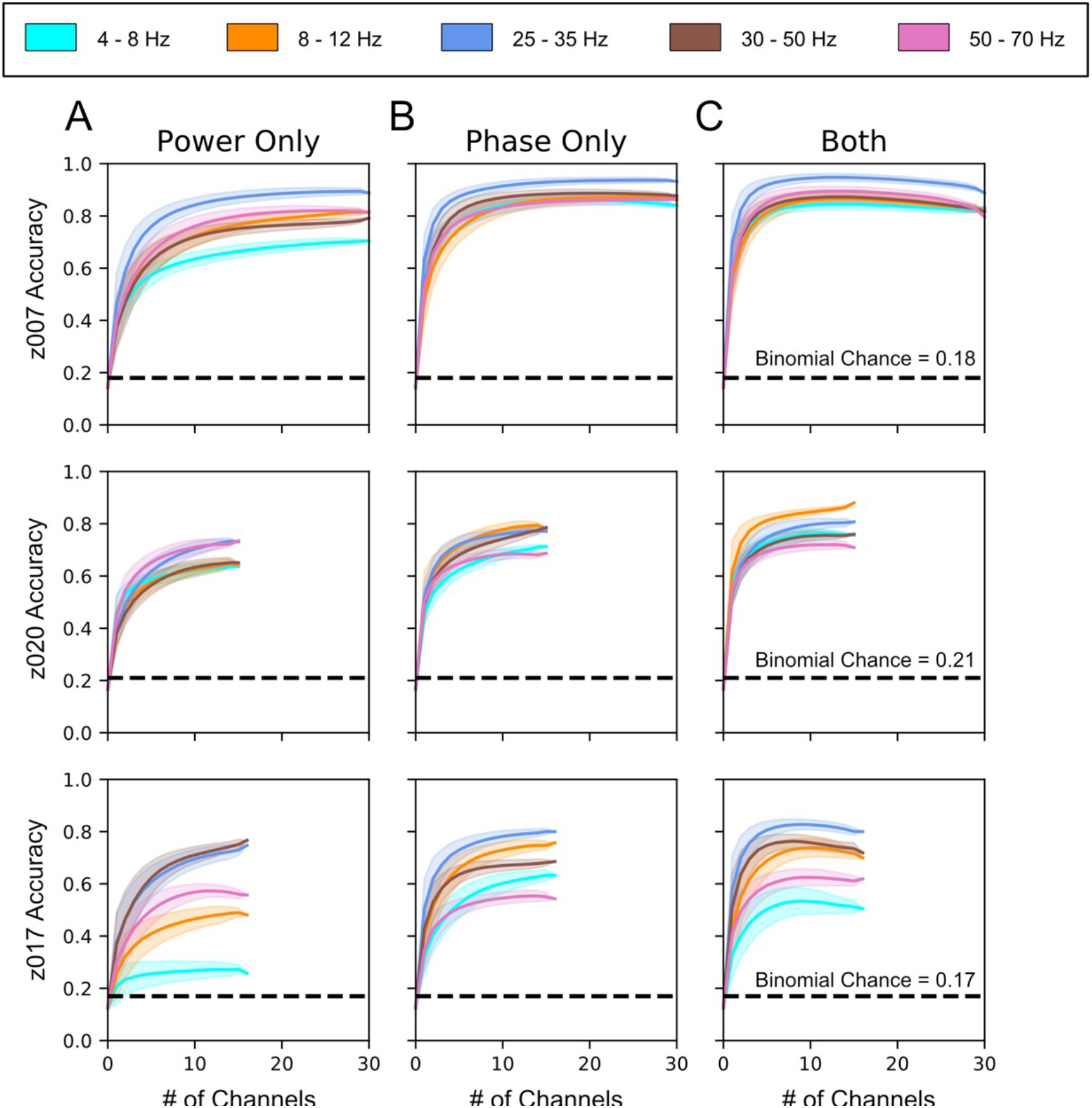
Both Phase and Power has independent/additive Information to classify syllable identity. Channel adding curves calculated by repeatedly training classifiers with an increasing number of randomly selected channels (see Methods). **Channel Dropping Curves of classifier performances with either (A) all information about phase removed, (B) all information about power removed, or (C) with both phase and power used as independent features. Each row corresponds to data from the highest yield day for each bird. z007 n=98 for each class n=7 (1, 2, 3, 4, 5, i, Silence), z020 n=91 for each class n = 6 (1, 2, 3, 4, i,** Silence), and for z017 n=52* for each class n=8 (1, 2, 3, 4, 5, 6, 7, Silence). Error bars represent the standard deviation over the bootstrapped analysis using n=5,000 repetitions across 5 cross-validation folds. The p-value for all of the Binomial chances calculated for each bird was 0.05. *The number of instances for each class was limited by Syllable 7, which is a intra-motif note.

**S13 Fig:**
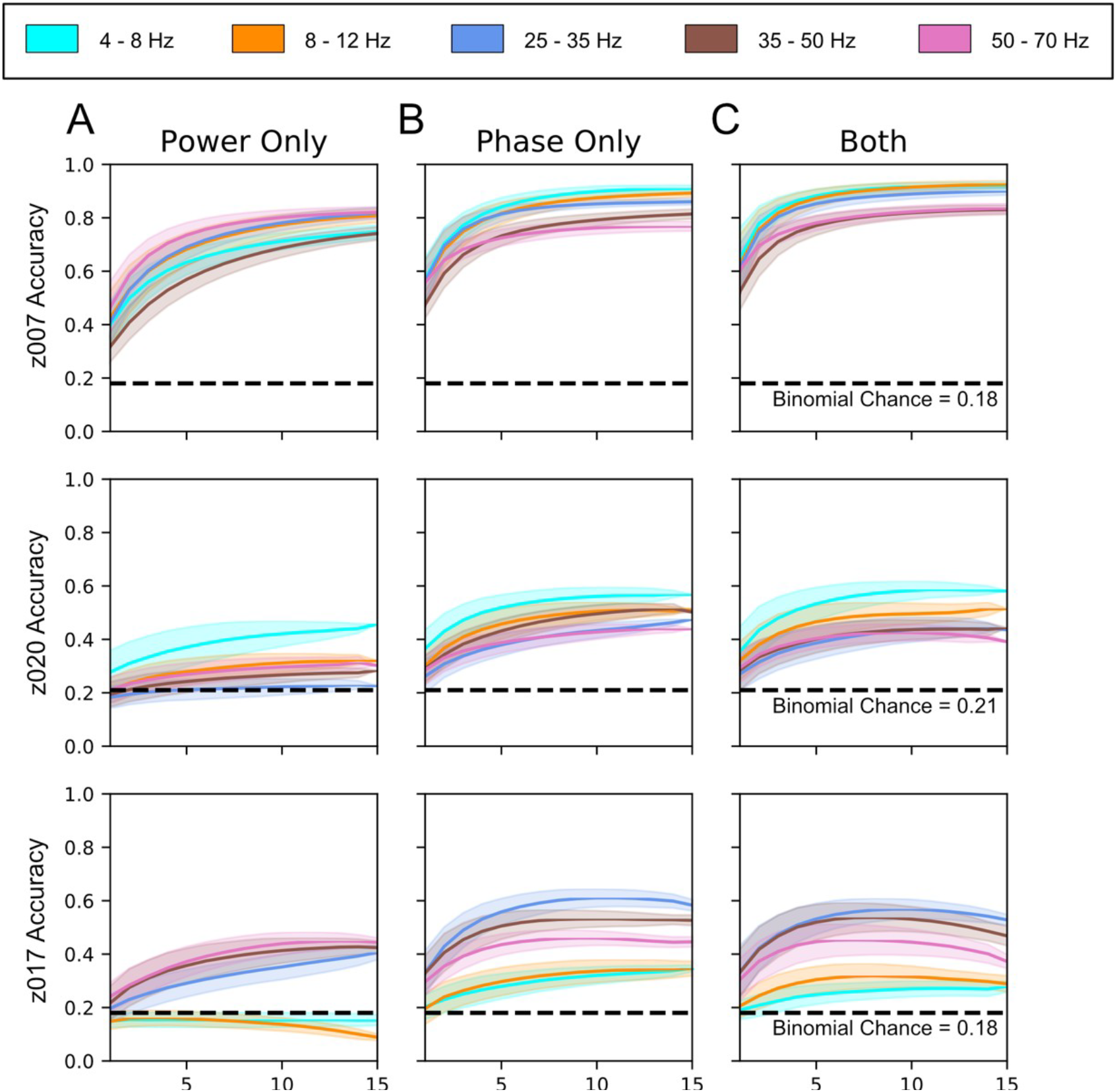
Channel Dropping Curves for the second highest yield days across subjects. Channel adding curves calculated by repeatedly training classifiers with an increasing number of randomly selected channels (see Methods). Channel Dropping Curves of classifier performances with either (A) all phase related information removed, (B) all power related information removed, or (C) with both phase and power used as independent features. Each row corresponds to data from the second highest yielding day for each bird. z007 n=71 for each class n=7 (1, 2, 3, 4, 5, i, Silence), z020 n=75 for each class n = 6 (1, 2, 3, 4, i, Silence), and for z017 n=41* for each class n=8 (1, 2, 3, 4, 5, 6, 7, Silence). Error bars represent the standard deviation over the bootstrapped analysis using n=5,000 repetitions across 5 cross-validation folds. The p-value for all of the Binomial chances calculated for each bird was 0.05. *The number of instances for each class was limited by Syllable 7, which is a intra-motif note.

**S14 Fig:**
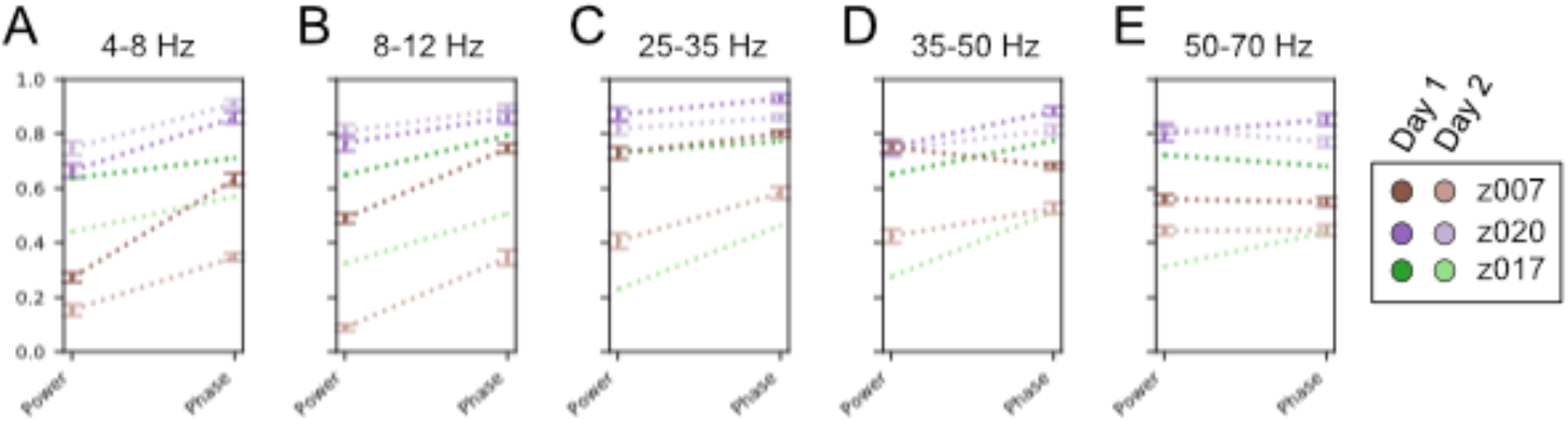
Difference in Decoding Accuracy between Phase Only and Power Only Classification. Classification Accuracy for each high yield day for each bird with 15 channels of neural data when all phase related information is removed, left, and all power related information is removed, right, for (A) 4-8 Hz Band, (B) 8-12 Hz Band, (C) 25-35 Hz Band, (D) 35-50 Hz Band, and € 50-70 Hz Band.

**S15 Fig:**
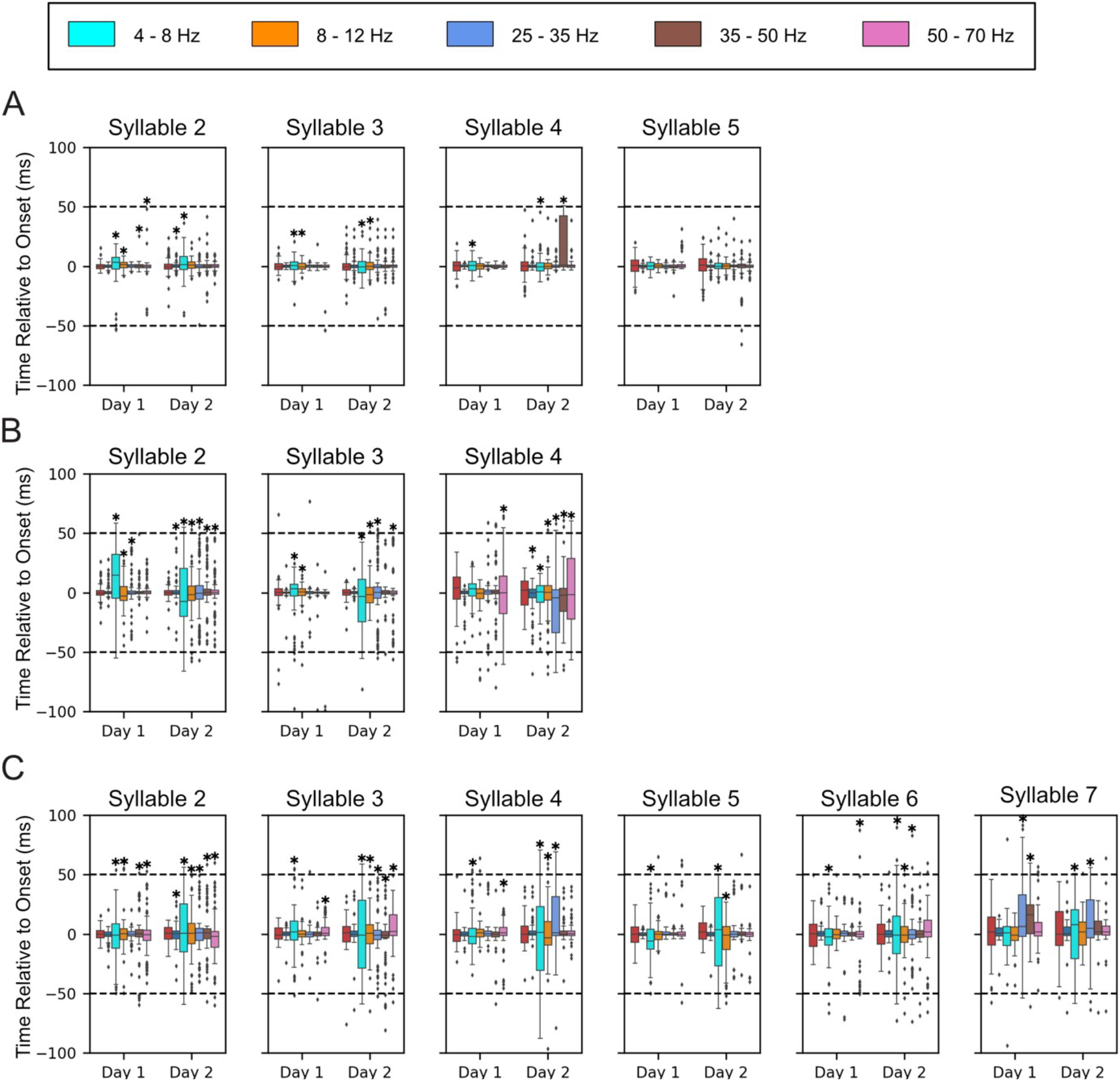
Onset detection across all high yield days. Boxplots of onset prediction times relative to the labeled onset time for each bird for every syllable for two highest yielding days. Each Column reflects the result for syllable number within the motif. Each row is for a specific bird with (A) corresponding to z007, (B) corresponding to z020, and (C) corresponding to z017. The order of each feature used is the same, going left to right: first is the stereotyped onset time using only the deterministic behavior, next is the results using all of the neural features, then each frequency band only in order from least to greatest (4-8 Hz, 8-12 Hz, 25-35 Hz, 35-50 Hz, and finally 50-70 Hz). The recording day designation number refers to the chronological order that the recordings took place. Statistical significance was calculated using the one sided Wilcoxon signed-rank test, and * denotes results that were not statistically significant when using the Benjamini-Hochberg False Discovery Rate. All other results p<0.05 and q<0.05.

## References

1. Goller F, Cooper BG. Peripheral Motor Dynamics of Song Production in the Zebra Finch. Ann N Y Acad Sci. 2004;1016: 130–152. doi:10.1196/annals.1298.009

2. Jarvis ED. Evolution of vocal learning and spoken language. Science. 2019;366: 50–54. doi:10.1126/science.aax0287

3. Nowicki S. Vocal tract resonances in oscine bird sound production: evidence from birdsongs in a helium atmosphere. Nature. 1987;325: 53–55. doi:10.1038/325053a0

4. Churchland MM, Cunningham JP, Kaufman MT, Ryu SI, Shenoy KV. Cortical Preparatory Activity: Representation of Movement or First Cog in a Dynamical Machine? Neuron. 2010;68: 387–400. doi:10.1016/j.neuron.2010.09.015

5. David A. Rosenbaum. Human Movement Initiation: Specification of Arm, Direction, and Extent. J Exp Psychol Gen. 1980;109: 444–474. doi:https://doi.org/10.1037/0096-3445.109.4.444

6. Fee MS, Kozhevnikov AA, Hahnloser RHR. Neural Mechanisms of Vocal Sequence Generation in the Songbird. Ann N Y Acad Sci. 2004;1016: 153–170. doi:10.1196/annals.1298.022

7. Ghez C, Hening W, Gordon J. Organization of voluntary movement. Curr Opin Neurobiol. 1991;1: 664–671. doi:10.1016/S0959-4388(05)80046-7

8. Kozhevnikov AA, Fee MS. Singing-Related Activity of Identified HVC Neurons in the Zebra Finch. J Neurophysiol. 2007;97: 4271–4283. doi:10.1152/jn.00952.2006

9. Daliparthi VK, Tachibana RO, Cooper BG, Hahnloser RH, Kojima S, Sober SJ, et al. Transitioning between preparatory and precisely sequenced neuronal activity in production of a skilled behavior. eLife. 2019;8: e43732. doi:10.7554/eLife.43732

10. Rajan R. Pre-Bout Neural Activity Changes in Premotor Nucleus HVC Correlate with Successful Initiation of Learned Song Sequence. J Neurosci. 2018;38: 5925–5938. doi:10.1523/JNEUROSCI.3003-17.2018

11. Bouchard KE, Mesgarani N, Johnson K, Chang EF. Functional organization of human sensorimotor cortex for speech articulation. Nature. 2013;495: 327–332. doi:10.1038/nature11911

12. Bouchard KE, Chang EF. Neural decoding of spoken vowels from human sensory-motor cortex with high-density electrocorticography. 2014 36th Annual International Conference of the IEEE Engineering in Medicine and Biology Society. Chicago, IL: IEEE; 2014. pp. 6782–6785. doi:10.1109/EMBC.2014.6945185

13. Martin S, Iturrate I, Millán J del R, Knight RT, Pasley BN. Decoding Inner Speech Using Electrocorticography: Progress and Challenges Toward a Speech Prosthesis. Front Neurosci. 2018;12: 422. doi:10.3389/fnins.2018.00422

14. Rabbani Q, Milsap G, Crone NE. The Potential for a Speech Brain–Computer Interface Using Chronic Electrocorticography. Neurotherapeutics. 2019;16: 144–165. doi:10.1007/s13311-018-00692-2

15. Sahin NT, Pinker S, Cash SS, Schomer D, Halgren E. Sequential Processing of Lexical, Grammatical, and Phonological Information Within Broca’s Area. Science. 2009;326: 445–449. doi:10.1126/science.1174481

16. Churchland MM, Santhanam G, Shenoy KV. Preparatory Activity in Premotor and Motor Cortex Reflects the Speed of the Upcoming Reach. J Neurophysiol. 2006;96: 3130–3146. doi:10.1152/jn.00307.2006

17. Tanji J, Evarts EV. Anticipatory activity of motor cortex neurons in relation to direction of an intended movement. J Neurophysiol. 1976;39: 1062–1068. doi:10.1152/jn.1976.39.5.1062

18. Svoboda K, Li N. Neural mechanisms of movement planning: motor cortex and beyond. Curr Opin Neurobiol. 2018;49: 33–41. doi:10.1016/j.conb.2017.10.023

19. Erlich JC, Bialek M, Brody CD. A Cortical Substrate for Memory-Guided Orienting in the Rat. Neuron. 2011;72: 330–343. doi:10.1016/j.neuron.2011.07.010

20. Guo ZV, Li N, Huber D, Ophir E, Gutnisky D, Ting JT, et al. Flow of Cortical Activity Underlying a Tactile Decision in Mice. Neuron. 2014;81: 179–194. doi:10.1016/j.neuron.2013.10.020

21. Li N, Chen T-W, Guo ZV, Gerfen CR, Svoboda K. A motor cortex circuit for motor planning and movement. Nature. 2015;519: 51–56. doi:10.1038/nature14178

22. Sainburg T, Theilman B, Thielk M, Gentner TQ. Parallels in the sequential organization of birdsong and human speech. Nat Commun. 2019;10: 3636. doi:10.1038/s41467-019-11605-y

23. Angrick M, Herff C, Mugler E, Tate MC, Slutzky MW, Krusienski DJ, et al. Speech synthesis from ECoG using densely connected 3D convolutional neural networks. J Neural Eng. 2019;16: 036019. doi:10.1088/1741-2552/ab0c59

24. Anumanchipalli GK, Chartier J, Chang EF. Speech synthesis from neural decoding of spoken sentences. Nature. 2019;568: 493–498. doi:10.1038/s41586-019-1119-1

25. Herff C, Schultz T. Automatic Speech Recognition from Neural Signals: A Focused Review. Front Neurosci. 2016;10. doi:10.3389/fnins.2016.00429

26. Stavisky SD, Willett FR, Wilson GH, Murphy BA, Rezaii P, Avansino DT, et al. Neural ensemble dynamics in dorsal motor cortex during speech in people with paralysis. eLife. 2019;8: e46015. doi:10.7554/eLife.46015

27. Brainard MS, Doupe AJ. What songbirds teach us about learning. Nature. 2002;417: 351–358. doi:10.1038/417351a

28. Nottebohm F. The Neural Basis of Birdsong. PLoS Biol. 2005;3: e164. doi:10.1371/journal.pbio.0030164

29. Okubo TS, Mackevicius EL, Payne HL, Lynch GF, Fee MS. Growth and splitting of neural sequences in songbird vocal development. Nature. 2015;528: 352–357. doi:10.1038/nature15741

30. Woolley SMN. Early experience shapes vocal neural coding and perception in songbirds. Dev Psychobiol. 2012;54: 612–631. doi:10.1002/dev.21014

31. Liberti WA, Markowitz JE, Perkins LN, Liberti DC, Leman DP, Guitchounts G, et al. Unstable neurons underlie a stable learned behavior. Nat Neurosci. 2016;19: 1665–1671. doi:10.1038/nn.4405

32. Poole B, Markowitz JE, Gardner TJ. The Song Must Go On: Resilience of the Songbird Vocal Motor Pathway. Coleman MJ, editor. PLoS ONE. 2012;7: e38173. doi:10.1371/journal.pone.0038173

33. Markowitz JE, Liberti WA, Guitchounts G, Velho T, Lois C, Gardner TJ. Mesoscopic Patterns of Neural Activity Support Songbird Cortical Sequences. Ashe J, editor. PLOS Biol. 2015;13: e1002158. doi:10.1371/journal.pbio.1002158

34. Picardo MA, Merel J, Katlowitz KA, Vallentin D, Okobi DE, Benezra SE, et al. Population-Level Representation of a Temporal Sequence Underlying Song Production in the Zebra Finch. Neuron. 2016;90: 866–876. doi:10.1016/j.neuron.2016.02.016

35. Schmidt MF. Pattern of Interhemispheric Synchronization in HVc During Singing Correlates With Key Transitions in the Song Pattern. J Neurophysiol. 2003;90: 3931–3949. doi:10.1152/jn.00003.2003

36. Bottjer SW, Johnson F. Circuits, hormones, and learning: Vocal behavior in songbirds. : 17.

37. Luo M, Perkel DJ. A GABAergic, Strongly Inhibitory Projection to a Thalamic Nucleus in the Zebra Finch Song System. J Neurosci. 1999;19: 6700–6711. doi:10.1523/JNEUROSCI.19-15-06700.1999

38. Nottebohm F, Stokes TM, Leonard CM. Central control of song in the canary, Serinus canarius. J Comp Neurol. 1976;165: 457–486. doi:10.1002/cne.901650405

39. Vates GE, Vicario DS, Nottebohm F. Reafferent thalamo-“cortical” loops in the song system of oscine songbirds. : 16.

40. Doupe AJ, Kuhl PK. BIRDSONG AND HUMAN SPEECH: Common Themes and Mechanisms. Annu Rev Neurosci. 1999;22: 567–631. doi:10.1146/annurev.neuro.22.1.567

41. Pfenning AR, Hara E, Whitney O, Rivas MV, Wang R, Roulhac PL, et al. Convergent transcriptional specializations in the brains of humans and song-learning birds. Science. 2014;346: 1256846–1256846. doi:10.1126/science.1256846

42. Buzsáki G, Anastassiou CA, Koch C. The origin of extracellular fields and currents — EEG, ECoG, LFP and spikes. Nat Rev Neurosci. 2012;13: 407–420. doi:10.1038/nrn3241

43. Carmena JM, Lebedev MA, Crist RE, O’Doherty JE, Santucci DM, Dimitrov DF, et al. Learning to Control a Brain–Machine Interface for Reaching and Grasping by Primates. Idan Segev, editor. PLoS Biol. 2003;1: e42. doi:10.1371/journal.pbio.0000042

44. Fetz EE. Operant Conditioning of Cortical Unit Activity. Science. 1969;163: 955–958. doi:10.1126/science.163.3870.955

45. Gilja V, Nuyujukian P, Chestek CA, Cunningham JP, Yu BM, Fan JM, et al. A high-performance neural prosthesis enabled by control algorithm design. Nat Neurosci. 2012;15: 1752–1757. doi:10.1038/nn.3265

46. Kubánek J, Miller KJ, Ojemann JG, Wolpaw JR, Schalk G. Decoding flexion of individual fingers using electrocorticographic signals in humans. J Neural Eng. 2009;6: 066001. doi:10.1088/1741-2560/6/6/066001

47. Miller KJ, Leuthardt EC, Schalk G, Rao RPN, Anderson NR, Moran DW, et al. Spectral Changes in Cortical Surface Potentials during Motor Movement. J Neurosci. 2007;27: 2424–2432. doi:10.1523/JNEUROSCI.3886-06.2007

48. Kosche G, Vallentin D, Long MA. Interplay of Inhibition and Excitation Shapes a Premotor Neural Sequence. J Neurosci. 2015;35: 1217–1227. doi:10.1523/JNEUROSCI.4346-14.2015

49. Leonardo A. Ensemble Coding of Vocal Control in Birdsong. J Neurosci. 2005;25: 652–661. doi:10.1523/JNEUROSCI.3036-04.2005

50. Miller KJ, Zanos S, Fetz EE, den Nijs M, Ojemann JG. Decoupling the Cortical Power Spectrum Reveals Real-Time Representation of Individual Finger Movements in Humans. J Neurosci. 2009;29: 3132–3137. doi:10.1523/JNEUROSCI.5506-08.2009

51. Brovelli A, Lachaux J-P, Kahane P, Boussaoud D. High gamma frequency oscillatory activity dissociates attention from intention in the human premotor cortex. NeuroImage. 2005;28: 154–164. doi:10.1016/j.neuroimage.2005.05.045

52. Canolty RT, Edwards E, Dalal SS, Soltani M, Nagarajan SS, Kirsch HE, et al. High Gamma Power Is Phase-Locked to Theta Oscillations in Human Neocortex. Science. 2006;313: 1626–1628. doi:10.1126/science.1128115

53. Pfurtscheller G, Graimann B, Huggins JE, Levine SP, Schuh LA. Spatiotemporal patterns of beta desynchronization and gamma synchronization in corticographic data during self-paced movement. Clin Neurophysiol. 2003;114: 1226–1236. doi:10.1016/S1388-2457(03)00067-1

54. Buzsáki G. Rhythms of the Brain. Oxford University Press; 2006. doi:10.1093/acprof:oso/9780195301069.001.0001

55. Jiang W, Pailla T, Dichter B, Chang EF, Gilja V. Decoding speech using the timing of neural signal modulation. 2016 38th Annual International Conference of the IEEE Engineering in Medicine and Biology Society (EMBC). Orlando, FL, USA: IEEE; 2016. pp. 1532–1535. doi:10.1109/EMBC.2016.7591002

56. Lewandowski BC, Schmidt M. Short Bouts of Vocalization Induce Long-Lasting Fast Gamma Oscillations in a Sensorimotor Nucleus. J Neurosci. 2011;31: 13936–13948. doi:10.1523/JNEUROSCI.6809-10.2011

57. Hyland Bruno J, Tchernichovski O. Regularities in zebra finch song beyond the repeated motif. Behav Processes. 2019;163: 53–59. doi:10.1016/j.beproc.2017.11.001

58. Miller KJ, Hermes D, Honey CJ, Hebb AO, Ramsey NF, Knight RT, et al. Human Motor Cortical Activity Is Selectively Phase-Entrained on Underlying Rhythms. Behrens T, editor. PLoS Comput Biol. 2012;8: e1002655. doi:10.1371/journal.pcbi.1002655

59. G. Makin J, Edward F. Chang, A. Moses D. Machine translation of cortical activity to text with an encoder-decoder framework. : 22.

60. Ezequiel Arneodo. Software and hardware designs for chronic, high channel count electrophysiology. Biocircuits Institute; 2016. Available: https://github.com/singingfinch/bernardo.git

61. Arneodo EM, Chen S, Gilja V, Gentner TQ. A neural decoder for learned vocal behavior. Neuroscience; 2017 Sep. doi:10.1101/193987

62. Gramfort A. MEG and EEG data analysis with MNE-Python. Front Neurosci. 2013;7. doi:10.3389/fnins.2013.00267

63. Paul Boersma, David Weenink. Praat: doing phonetics by computer. 2018. Available: http://www.praat.org/

64. Ludwig KA, Miriani RM, Langhals NB, Joseph MD, Anderson DJ, Kipke DR. Using a Common Average Reference to Improve Cortical Neuron Recordings From Microelectrode Arrays. J Neurophysiol. 2009;101: 1679–1689. doi:10.1152/jn.90989.2008

65. Delorme A, Makeig S. EEGLAB: an open source toolbox for analysis of single-trial EEG dynamics including independent component analysis. J Neurosci Methods. 2004;134: 9–21. doi:10.1016/j.jneumeth.2003.10.009

66. Berens P. CircStat: A MATLAB Toolbox for Circular Statistics. J Stat Softw. 2009;31. doi:10.18637/jss.v031.i10

67. Fisher NI. Statistical Analysis of Circular Data. 1st ed. Cambridge University Press; 1993. doi:10.1017/CBO9780511564345

68. Jammalamadaka SR, Sengupta A. Topics in circular statistics. River Edge, N.J: World Scientific; 2001.

69. Zar JH. Biostatistical analysis. 5th ed. Upper Saddle River, N.J: Prentice-Hall/Pearson; 2010.

70. Pedregosa F, Varoquaux G, Gramfort A, Michel V, Thirion B, Grisel O, et al. Scikit-learn: Machine Learning in Python. Mach Learn PYTHON. 2011; 6.

